# Bioluminescent Synthetic Cells Communicate with Natural Cells and Self-Activate Light-Responsive Proteins

**DOI:** 10.1101/2021.05.20.444896

**Authors:** Omer Adir, Ravit Abel, Mia R. Albalak, Lucien E. Weiss, Gal Chen, Amit Gruber, Oskar Staufer, Jeny Shklover, Janna Shainsky-Roitman, Ilia Platzman, Lior Gepstein, Yoav Shechtman, Benjamin A. Horwitz, Avi Schroeder

## Abstract

Development of regulated cellular processes and signaling methods in synthetic cells is essential for their integration with living materials. Light is an attractive tool to achieve this, but the limited penetration depth into tissue of visible light restricts its usability for *in-vivo* applications. Here, we describe the synthesis and application of blue-light-generating synthetic cells using bioluminescence, dismissing the need for an external light source. First, the lipid membrane and internal composition of light-producing synthetic cells were optimized to enable high-intensity emission. Next, we show these cells’ capacity for triggering bioprocesses in natural cells by initiating asexual sporulation of dark-grown mycelial cells of the fungus *Trichoderma atroviride* in a quorum-sensing like mechanism. Finally, we demonstrate regulated transcription and membrane recruitment in synthetic cells using bioluminescent self-activating fusion proteins. These functionalities pave the way for deploying synthetic cells as embeddable microscale light sources that are capable of activating engineered processes inside tissues.

---

In recent years, there has been a growing interest in the field of synthetic cells for therapeutics, diagnostics and research on the origin of life. These cell-mimicking microparticles, are designed from the bottom-up to reconstitute various processes of living cells and introduce entirely new functionalities that are not present naturally^1–3^. Biological processes such as protein expression, ATP production, DNA replication and cytoskeleton re-arrangement have all been successfully reconstructed in synthetic cells, yielding new insights in the isolated and controlled model environment^4–7^. Due to their tunable properties (i.e., size, composition, membrane rigidity) and engineerability, promising clinical applications in diagnostics and therapeutics are being realized as well. These include insulin-secreting synthetic beta cells and therapeutic-protein-producing synthetic cells that express pseudomonas exotoxin A (PE) to eliminate 4T1 breast cancer cells, both of which have already been tested in animal models *in-vivo*^8–9^. The implementation of increasingly complex functions in synthetic cell technologies requires inputs and outputs, i.e. cell signaling, that can be used orthogonally^10,11^. This is of particular importance for interfacing synthetic cells in living tissue.

Previous reports have demonstrated chemical communication between synthetic cells and bacterial cells as well as between synthetic cells^12,13^. For example, isopropyl b-D-1-thiogalactopyranoside (IPTG), arabinose and C6-HSL (N-(3-oxohexanoyl)-L-homoserine lactone) have been used as signaling molecules to control gene expression and differentiation in synthetic cells.^14,15^ Light has also been used to stimulate cellular processes in synthetic cells. UV radiation (365 nm) was used to initiate transcription and activate communication in synthetic cells and tissues by unlocking DNA photo-caging^16,17^. Another example was demonstrated using blue light to trigger ATP production in synthetic cells utilizing bacteriorhodopsin to form a proton gradient and drive ATP synthase activity^18^. This wide diversity of light-responsive elements that are activated by light from different parts of the spectrum provides multichannel signaling opportunities. Problematically, using light over much of the visible spectrum for *in-vivo* applications often requires invasive transplantations of external light sources^19,20^. A translational alternative for using external light sources is to generate light from synthetic cells localized in the target area. This can be achieved by exploiting bioluminescent reactions catalyzed by enzymes from the luciferase family, which have been widely used in research for biological reporter assays and *in-vivo* imaging^21,22^.

Here, we engineer bioluminescent signaling mechanisms in synthetic cells for the purpose of activating cellular processes in both natural and synthetic cells. These synthetic cells were composed of giant unilamellar vesicles (GUVs) encapsulating a bacterial-based, cell-free protein synthesis (CFPS) system (Fig. 1a). Light-generating synthetic cells expressing *Gaussia* luciferase (Gluc) were first designed and optimized for emission intensity. Their capacity to activate biological processes was demonstrated by initiating photoconidiation in fungal cells. For biological processes involving light-responsive proteins that required higher intensities, novel self-activating fusion proteins were engineered by coupling Gluc with the responder protein, facilitating activation by bioluminescence resonance energy transfer (BRET). This was used to activate transcription in synthetic cells using a fusion protein of Gluc and the light-activated transcription factor EL222. Finally, we engineered a membrane recruitment mechanism in synthetic cells using a membrane-bound fusion protein of Gluc and iLID, that dimerized with a sspB-tagged protein when the bioluminescent reaction was initiated. These new synthetic cell functionalities present opportunities for activation and control of synthetic and natural cells alike.

**Figure 1.**
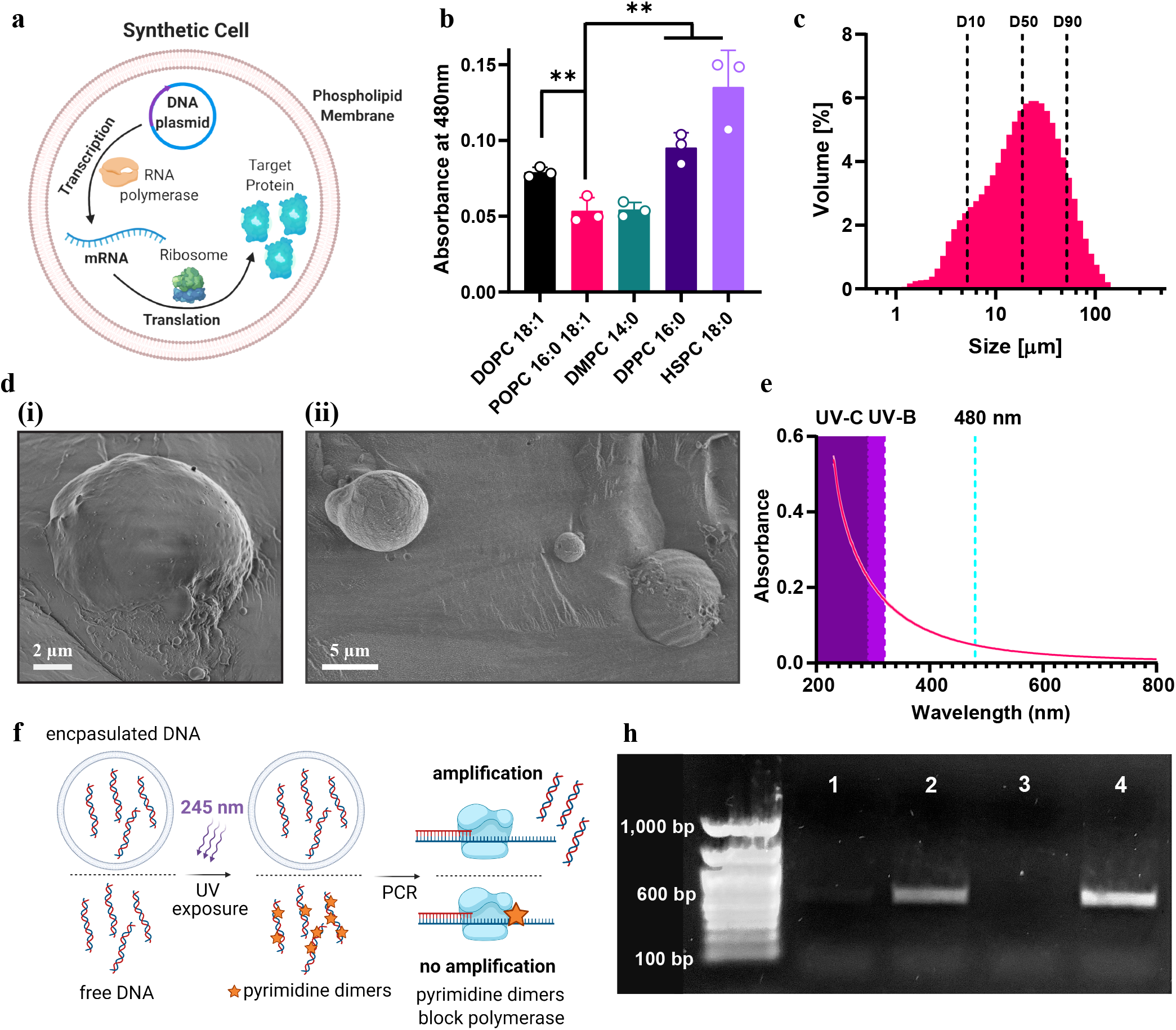
Optimizing the lipid composition for light-interacting synthetic cells. **a,** Illustration of a protein producing synthetic cell. **b,** The effect of the lipid membrane composition on blue light absorbance in liposomes. Data is expressed as the mean ± standard deviation (n=3 independent samples). Nested two-tailed t-test P values; **P ≤ 0.0079. **c,** Size distribution of POPC:cholesterol synthetic cells measured with light scattering. Data is expressed as mean of n=3 independent samples. **d,** Morphology of POPC:cholesterol synthetic cells imaged with cryogenic scanning electron microscopy (cryo-SEM). **e,** The Absorbance spectrum of POPC liposomes between 230 and 800 nm. **f,** Illustration of a linear DNA oligonucleotide, encapsulated in a synthetic cell or free in solution, exposed to UV radiation. Formation of pyrimidine dimers is detected using PCR amplification. **h,** Gel electrophoresis of the amplified PCR product of linear DNA after exposure to UV radiation with and without encapsulation in synthetic cells. Lane 1: free DNA exposed to UV. Lane 2: encapsulated DNA exposed to UV. Lane 3: no DNA control. Lane 4: untreated DNA control.

## Results

### Optimizing the lipid composition of light-interacting synthetic cells

The phospholipid membrane is the main physical barrier for light emission or light absorption in synthetic cells composed of GUVs, and must therefore have favorable optical properties. Hence, the first step in constructing light-generating and light-responding synthetic cells is to optimize their membrane composition. In this work, we focused on interactions with blue light (480 nm), which can be used to activate photoreceptors such as light-oxygen-voltage-sensing (LOV) domains present in many photoactivatable proteins, and is emitted by certain types of luciferases, including *Renilla* and *Gaussia* luciferase^23,24^. Furthermore, the use of blue light for *in-vivo* applications traditionally poses a challenge for non-implanted sources due to its poor tissue penetration.

Factors affecting light transmission through the lipid membrane were investigated and the phospholipid composition of the membrane was altered accordingly to minimize attenuation. First, the absorbance of 480 nm light by 100-nm liposomes composed of a single phospholipid type was measured (Fig. 1b). This liposome size was selected for this measurement due to its lower polydispersity index (Fig. S1), thereby reducing sample-to-sample variation originating from size-dependent light interactions (i.e. light scattering). The lipid-light interactions demonstrated a positive correlation between the phospholipid tail length and light absorbance (Fig 1b). L-α-phosphatidylcholine, hydrogenated (Soy), (HSPC) which is composed of 88.6% of 18:0 fatty acids had the highest absorbance, which was more than 1.5-times higher than 1,2-dipalmitoyl-sn-glycero-3-phosphocholine (DPPC) with 16:0 tails and 2.6-times higher than 1,2-dimyristoyl-sn-glycero-3-phosphocholine (DMPC) with 14:0 tails. This trend was also apparent in liposomes composed of unsaturated phospholipids, in which the 18:1 lipid 1,2-dioleoyl-sn-glycero-3-phosphocholine (DOPC) had higher light attenuation compared to 16:0,18:1 lipid 1-palmitoyl-2-oleoyl-glycero-3-phosphocholine (POPC). Moreover, liposomes composed of unsaturated lipids had lower absorbance relative to those made of saturated-lipids with similar tail lengths (more than 1.7-times difference between the absorbance of HSPC and DOPC).

While both DMPC and POPC liposomes had low light absorbance, the latter was selected as the main lipid in the synthetic cell due to its lower melting temperature (Tm) of −2°C which enabled it to remain in a liquid phase during the synthetic cell production process performed at 4°C. Although cholesterol does contribute to the light absorbance (Fig. S2), its presence in the synthetic cell membrane (50% w/w ratio) is crucial to enhance membrane stability, which is important as the incubation for protein production is performed at 37 °C.

Once the lipid composition of the membrane was selected, synthetic cells were produced using the emulsion transfer method. This method’s simplicity, high encapsulation efficiency, and high yield make it preferable for achieving high overall protein production^25^. Light scattering was used to determine the obtained size distribution, spanning 1-200 μm with average D10, D50 and D90 values (corresponding to the 10%, 50% and 90% marks in the cumulative size distribution curve) of 5.2, 18.6 and 51.8 μm, respectively (Fig. 1c). Cell morphology and size range were further validated with cryogenic scanning electron microscopy (cryo-SEM, Fig. 1d, S3).

We then scanned the absorbance spectrum of different phospholipids at different wavelengths (230-800 nm) to understand if the characteristics recognized for 480 nm light were relevant for other wavelengths that could also be used for light signaling. While the particles exhibited little light attenuation at wavelengths longer than 480 nm, a sharp increase in the absorbance of all the liposomal formulations was observed in the UVC (200-290 nm) and UVB (290-320 nm) range (Fig. 1e, S4). Based on these results, we hypothesized that the phospholipid membrane could protect DNA from UV radiation damage and tested this using our synthetic cell platform (Fig. 1f-h).

Exposure of DNA to UV radiation is known to lead to the formation of pyrimidine dimers and other DNA photoproducts that damage the DNA functionality and lead to subsequent biological effects (Fig. 1f)^26,27^. We exposed a 600-bp linear DNA oligonucleotide to UV radiation for 20 minutes before or after its encapsulation in a synthetic cell and detected the formation of DNA lesions with polymerase chain-reactions (PCR), which is inhibited by these lesions^28^. Encapsulation of the DNA in synthetic cell yielded evident PCR amplification indicating improved UV protection in comparison to the DNA that was irradiated prior to its encapsulation and yielded almost no PCR product at all. In addition to their eminent role as compartmentalizing structure, this highlights the importance of phospholipid membranes as UV-protecting scaffolds.

### Engineering light-generating synthetic cells

Following the optimization of the synthetic cells’ membrane, we engineered synthetic cells capable of generating light by expressing luciferase for catalyzing a photon-emitting enzymatic reaction. Considering energy consumption, emitted wavelength, and total light emission intensity parameters, we chose to focus on two luciferase types, *Renilla* luciferase (Rluc) and a mutated variant of *Gaussia* luciferase (M43L, M110L)^29^ sourced from the organisms *Renilla reniformis* and *Gaussia princeps* respectively. Both types of luciferase catalyze an ATP-independent bioluminescent oxidation reaction of the substrate coelenterazine, conserving energy for the synthetic cell^24,30^. In terms of emission wavelength, both enzymes generate photons in the blue-range, suitable for LOV domain activation. Nevertheless, while Rluc has rapid flash kinetics, the Gluc variant exhibits brighter and longer half-life of illumination (glow-like kinetics)^29^.

Initial comparison of light production of both luciferase types in CFPS reactions with *E. coli BL21(DE3)* lysates indicated 2500-fold higher light emission in Rluc-expressing synthetic cells in comparison to weak illumination produced from the Gluc-expressing synthetic cells (Fig. 2a). Most likely, the poor production of Gluc in the synthetic cells was due to misfolding of its five disulfide bonds in the reducing environment of the synthetic cells that contains both 1,4-Dithiothreitol (DTT) and 2-mercaptoethanol^31^. Moreover, oxidative folding for disulfide bond generation in *E. coli* is performed in the periplasm and not in the reducing environment of the bacterial cytoplasm which is a main component of the lysate^32^. Therefore, the synthetic cells’ internal composition was redesigned. Reducing agents DTT and 2-mercaptoethanol were excluded from the internal solution, and glutathione and disulfide bond isomerase C (DsbC) from *E. coli* were added to improve Gluc folding^33^. A range of oxidized and reduced glutathione concentrations were added to the inner solution of the synthetic cells and light emission after CTZ addition was measured (Fig. S5). 4 mM of oxidized and 1 mM of reduced glutathione were found to be optimal for Gluc production with the *BL21(DE3)* lysate. After further addition of 75 μg ml^−1^ of DsbC (Fig. S6) to the glutathione-supplemented synthetic cell internal solution, light emission from Gluc-expressing synthetic cells was more than 100-fold higher than that of the Rluc-expressing synthetic cells (Fig. 2b), and was easily visible by eye in solution (Fig. 2c). Gluc expression in synthetic cells was quantified with western blot analysis demonstrating production of 43 ng μl^−1^ (Fig. 2d) of protein. This translates to an average of 25 pg of Gluc per synthetic cell, considering a concentration of 1.68·10^6^ synthetic cells ml^−1^, as quantified using multispectral imaging flow cytometry.

**Figure 2.**
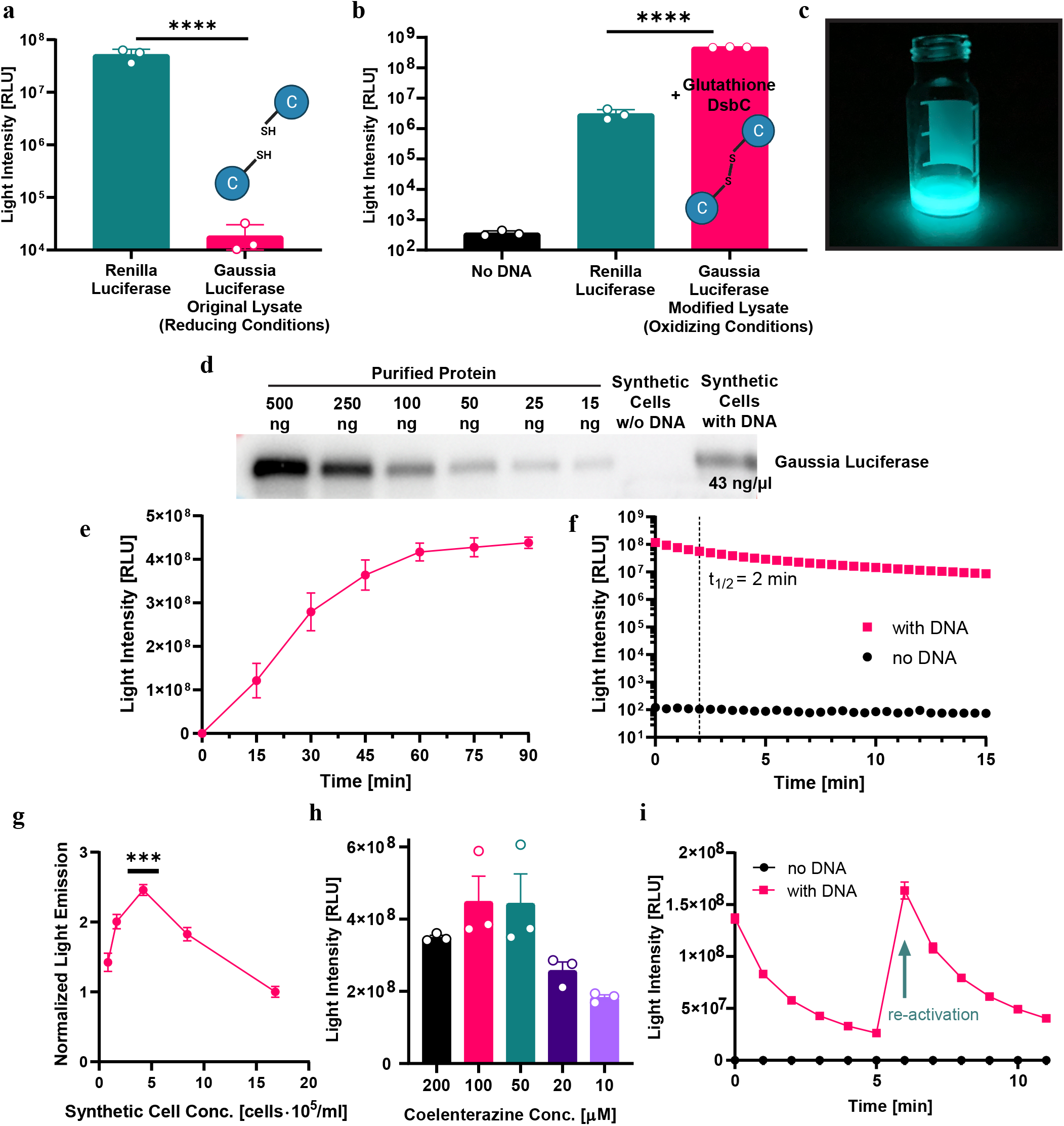
Engineering light-generating synthetic cells. **a,** Comparison of light emission in cell-free protein synthesis reactions expressing *Renilla* Luciferase (Rluc) or *Gaussia* luciferase (Gluc) using unmodified lysate (reducing conditions). Nested two-tailed t-test P value; ****P < 0.0001. **b,** Light emission in synthetic cells expressing Gluc using modified lysate with addition of glutathione and disulfide bond isomerase C (DsbC) to produce an oxidizing environment in the synthetic cells, compared to light emission in Rluc-expressing synthetic cells. Nested one-way ANOVA with multiple comparisons test adjusted P value; ****P<0.0001. **c,** Light emission from a Gluc-expressing synthetic cell solution. **d,** Western blot quantification of Gluc production in synthetic cells. **e,** Gluc production kinetics in synthetic cells at 37°C. **f,** Kinetics of the Gluc enzymatic reaction in synthetic cells after one addition of 100 μM coelenterazine. **g,** Light emission from Gluc-expressing synthetic cells diluted to different concentrations after incubation. Nested one-way ANOVA with multiple comparisons test adjusted P value; ***P ≤ 0.0006. **h,** Light emission in ranging coelenterazine concentrations in Gluc-expressing synthetic cells. Data is expressed as a mean ± s.e.m. (n=3 independent samples). Ordinary one-way ANOVA P value; *P = 0.013. **i,** Temporal control over light emission in Gluc-expressing synthetic cells with two 0.125 nmol coelenterazine additions (second addition marked with a green arrow). for a,b,e,f,g,i Data is expressed as a mean ± standard deviation (n=3 independent samples).

The protein production and light-emission properties of Gluc-expressing synthetic cells were further characterized under different incubation times, cell and substrate concentrations. Protein-production kinetics in synthetic cells at 37 °C were monitored using light-emission assays performed at different incubation times (Fig. 2e). Gluc levels increased over the first 60 minutes and plateaued at 90 minutes. Light emission kinetic measurements demonstrated a t1/2 of two minutes for synthetic cells incubated with 100 μM of CTZ (Fig. 2f). After 15 minutes, the illumination intensity was approximately 7% of the initial intensity measured and still more than 100,000 fold higher than the intensity measured from control synthetic cells without a DNA template encoding for luciferase.

Next, the dependency of light intensity on the synthetic cell concentration was demonstrated (Fig. 2g). A concentration of 420,000 synthetic cells ml^−1^ was found to generate the maximal light intensity upon CTZ addition and was used for the subsequent experiments. In comparison, 2 and 4-times higher cell concentrations and 2.5 and 5-times lower cell concentrations had significantly lower light emission. Increasing concentrations of synthetic cells increase the total Gluc concentration, but also the total phospholipid and cholesterol concentration which attenuate light emission from the solution. The yield of emitted light is thus balanced between these two parameters.

Light intensity was also controlled by altering the substrate concentration, demonstrating an increase in the generated signal between 10 μM and 100 μM CTZ, and a slight decrease for 200 μM CTZ (Fig. 2h). Lower light intensities were also detectable using lower concentrations of substrate, down to 10 nM of CTZ (Fig. S7). Temporal control of illumination was achieved by timed addition of the substrate to the solution (0.125 nmol CTZ). The emission of synthetic cells was restored when an additional dose of 0.125 nmol of substrate was added. The light intensity generated upon additional doses of substrate following the decay of the initial light emission reached intensities similar to those produced in the first illumination pulse (Fig. 2i). This regeneration ability enables high light intensity levels to be maintained when prolonged illumination is required, and to perform multiple activations at different time points.

### Light-producing synthetic cells activate fungal cells

To demonstrate intercellular signaling between light-producing synthetic cells and light-responsive natural cells we employed the soil fungus *Trichoderma atroviride*. Two blue-light regulator (BLR) proteins control the photo-activation of this fungus in response to blue light, triggering several downstream processes including conidiation and the expression of the DNA repairing photolyase enzyme^34–36^. *T. atroviride* were plated and grown for 48 hours in dark room conditions to prevent activation by external light. 24 hours after a 1-minute exposure to blue LEDs (15 mW cm^−2^), a peripheral ring of spores was evident on the border of the fungi colony (Fig. S8). Alternatively, synthetic cell illumination was supplied in two consecutive induction rounds, each with 500 μL of synthetic cells supplemented with 100 μM CTZ that were added to a restricted section in the fungi plate and incubated for 15 minutes (Fig. 3a). After an incubation period of 24 hours in the dark following induction, conidiation was quantified by calculating the percentage of sporulated area out of the total area exposed to synthetic cells (Fig. 3b-c). Fungi incubated with Gluc-expressing synthetic cells demonstrated an average of 9.1% sporulated area, a 23-times increase in comparison to fungi incubated with synthetic cells without a DNA template (Fig. 3c).

**Figure 3.**
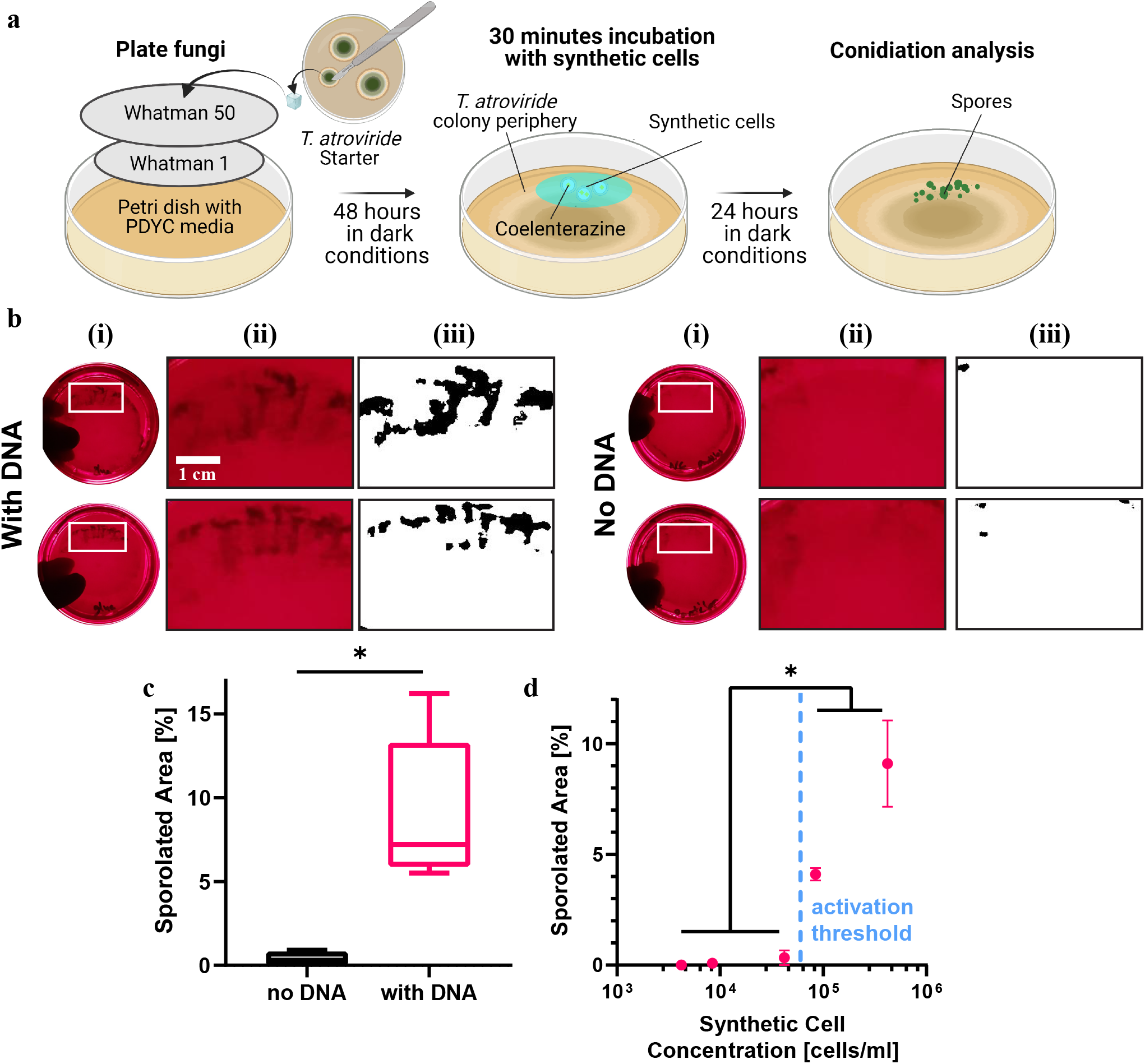
Activation of fungal cells using light-producing synthetic cells. **a,** Illustration of the experimental setup for photo-activation of conidiation in *Trichoderma atroviride* with Gluc-expressing synthetic cells. **b,** (i) Representative images of *Trichoderma atroviride* plates after exposure to Gluc-expressing synthetic cells or Synthetic cells without DNA. (ii) A magnified image and a (iii) black and white thresholded image of the plate area marked with a white rectangle in which the synthetic cells were localized. **c,** Quantitative analysis of the sporulated area out of the total area exposed to synthetic cells. Data is expressed as min-to-max box plot ± standard deviation (n=5 independent samples). Welch’s two-tailed t-test P value; *P = 0.0103. **d,** The effect of varying synthetic cell concentration on photo-activation of conidiation in *Trichoderma atroviride*. Data is expressed as a mean ± s.e.m. (between n=3 to n=5 independent samples). Welch’s two-tailed t-test P value; *P ≤ 0.01.

The effect of the synthetic cells’ concentration on the activation of the BLR pathway had quorum sensing characteristics. A range of synthetic cell dilutions (between 4,200 to 420,000 cells ml^−1^) were incubated on the *T. atroviride* plates and sporulation was quantified (Fig. 3d). A minimal concentration of 84,000 synthetic cells ml^−1^ was required for activation of conidiation, with a ~2.2-times increase in sporulated area observed between 84,000 cells ml^−1^ and 420,000 cells ml^−1^. In a previous study, Horwitz et al.^34^ showed that approximately 10^19^ photons m^−2^ of broadband blue light were necessary for initial activation of the *T. atroviride* culture in similar experimental conditions. This provides an approximate measure of the total photon flux produced by the synthetic cells during the experiment. The observed synthetic cell concentrationdependent response of *T. atroviride* resembles chemical quorum sensing mechanisms in other species, yet here the sensing is based on light dosage and can occur even when the synthetic and natural cells are separated by a physical barrier.

#### Bioluminescent self-activation of transcription in synthetic cells

Next, we demonstrate auto-activation of two different cellular processes in synthetic cells using bioluminescent reactions: induction of protein expression and membrane recruitment. To achieve this, we designed self-activating fusion proteins composed of Gluc and a light-responding domain that initiate these processes.

Control over protein expression was performed using the light-inducible bacterial transcription factor EL222, that dimerizes when exposed to blue light and binds to a specific region in the pBLind promoter to initiate transcription (Fig. 4a)^37,38^. To simplify the integration of EL222 into the synthetic cell inner solution, calibration of the required EL222 concentration was initially performed in CFPS reactions (before encapsulation in synthetic cells) using an external LED blue light source for activation. Rluc was used as a reporter protein, and a DNA plasmid containing its reading frame after the pBLind promoter was prepared. Purified EL222 was added in concentrations of 2.5 μM, 5 μM and 10 μM to the CFPS mix and incubated for 1 hour in dark or light conditions (Fig. 4b). Rluc expression levels were analyzed by adding CTZ and measuring luminescence. EL222 had the most significant light-to-dark ratio at 2.5 μM, and demonstrated significant light-to-dark differences at 5 and 10 μM. Production of monomeric RFP (mRFP1), that is characterized by a longer folding time, in CFPS reactions with 2.5 μM EL222 was subsequently tested (Fig. 4c). Elevated mRFP1 levels were evident in the reaction mix containing EL222 and mRPF1 DNA after 5 hours of incubation with blue-light, in comparison to the same solution incubated in dark conditions, and continued to increase through the 12 hours measured (Fig. 4c, pink and black filled circles). Some increase in signal was evident in the dark-incubated sample without DNA (Fig. 4c, hollow black circles), and is associated with increase in auto fluorescence over time.

**Figure 4.**
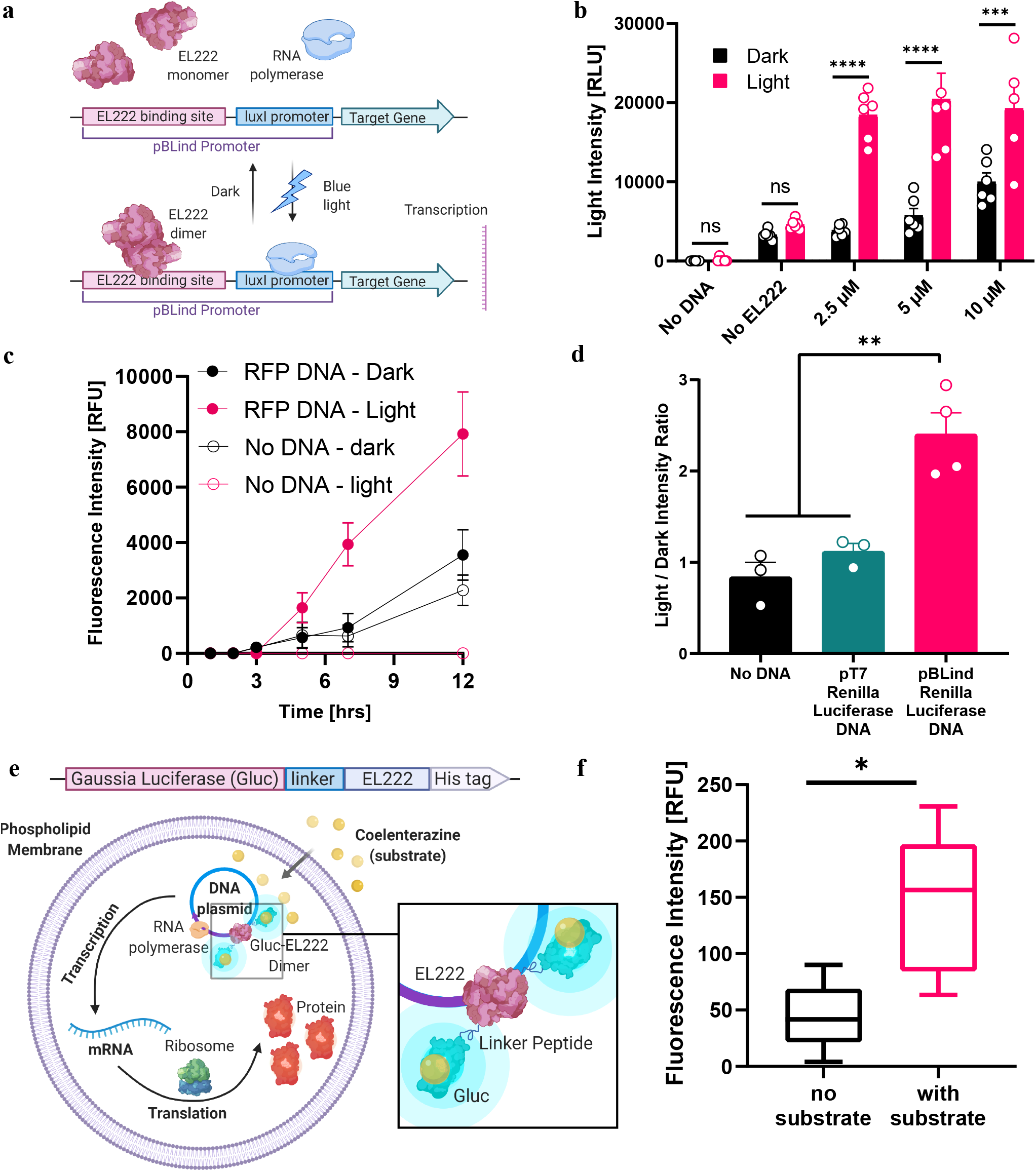
Bioluminescent signaling self-activates transcription in synthetic cells. **a,** Illustration of the light-dependent transcription mechanism mediated by the transcription factor EL222. **b,** EL222 concentration affected the production of *Renilla* luciferase (Rluc) in cell-free protein synthesis (CFPS) reactions under light and dark conditions. Data is expressed as a mean ± standard deviation (between n=5 to n=6 independent samples). Two-way ANOVA with multiple comparisons test adjusted P value; ***P = 0.0005; ****P < 0.0001. **c,** RFP production kinetics in CFPS reactions supplemented with EL222 under dark and light conditions, with or without RFP DNA. Data is expressed as a mean ± s.e.m. (between n=4 to n=6 independent samples). **d,** Light-to-dark ratio of Rluc expression in synthetic cells containing EL222 and a DNA plasmid expressing Rluc under different promoters. Data is expressed as a mean ± s.e.m. (between n=3 to n=4 independent samples). One-way ANOVA with multiple comparisons test adjusted P value; **P ≤ 0.005. **e,** A block diagram of the Gluc-EL222 fusion protein elements. Below, a schematic representation of a synthetic cells containing the Gluc-EL222 fusion protein for bioluminescent activation of transcription. **f,** RFP production in synthetic cells containing the Gluc-EL222 protein with or without addition of coelenterazine. Data is expressed as min-to-max box plot ± standard deviation (n=5 independent samples). Student’s two-tailed t-test P value; *P = 0.0134.

After establishing the reaction conditions in CFPS, we integrated the EL222 system in synthetic cells and activated expression of Rluc using an external LED light. The higher light absorbance of synthetic cells compared to CFPS required to increase the light intensity used for activation, while avoiding overheating that might lead to protein denaturation. Therefore, the light intensity was amplified from approximately 12 W m^−2^ to 19 W m^−2^ with on-off intervals of 20 and 70 seconds respectively. Under these conditions, synthetic cells encapsulating EL222 monomers and a DNA plasmid expressing Rluc under the pBLind promoter showed 2.4-fold increase in Rluc expression in light vs. dark conditions (Fig. 4d). In comparison, synthetic cells with EL222 and no DNA or a DNA plasmid with Rluc expressed under the viral T7 promoter which is not specific to EL222, showed negligible changes.

In order to activate EL222 with bioluminescence, close proximity between the light source and the responsive protein is required. For this purpose, we designed a new fusion protein, with Gluc on its N-terminal end connected through a flexible peptide linker to EL222 on its C-terminal end (Fig. S9). This construct utilizes BRET to exploit the energy from the bioluminescent luciferase reaction on its substrate CTZ, to directly activate the target protein (Fig. 4e). We added the Gluc-EL222 protein to the synthetic cell interior instead of the native EL222, and followed mRFP1 production with and without the addition of CTZ (Fig. 4f, S10). A 3.2-time increase in mRFP1 production was observed when CTZ was added to the synthetic cells compared to synthetic cells that were incubated in the dark without CTZ. This demonstrates the functionality of the fusion protein as a self-activating transcription factor in synthetic cells.

### Bioluminescent self-activated membrane recruitment in synthetic cells

To further explore the potential of bioluminescent self-activating processes in synthetic cells, we incorporated light-controlled protein recruitment capabilities to the synthetic cell membrane. Specifically, this was performed using a hetero-dimerization reaction with one monomer conjugated to the synthetic cell membrane (iLID) and the other free in solution (sspB-Nano, Fig. S11)^39^. We incorporated DGS-NTA-Ni lipids into the synthetic cell membrane to bind his-tagged iLID and monitored the localization of mRFP1 fused to the sspB-Nano protein. In these experiments, we prepared the synthetic cells with a droplet-based manual emulsification method, which is based on charge-mediated liposome fusion inside surfactant-stabilized droplets and enables higher flexibility in lipid selection in comparison to the emulsion-transfer method^40^. This allowed us to fabricate synthetic cells with a membrane composition of 73.5 mol% POPC, 20% DOPG, 5% DGS-NTA-Ni, 1% DOPE-PEG4-biotin and 0.5% DOPE-Cy5.

Gluc-expressing CFPS reactions did not provide sufficient light to activate the LOV domain of the iLID and initiate sspB-Nano recruitment to the synthetic cell membrane (Fig. S12). We therefore engineered a second fusion protein, N-terminal Gluc fused to C-terminal iLID with a linker peptide and an N-terminal his-tag, harnessing BRET again for efficient activation of the light-responsive domain (Fig. 5a, S13). This structure facilitates binding of the fusion protein to the synthetic cell membrane from the Gluc end and exposes the iLID to the extracellular environment with lower steric interference (Fig 5a). Gluc-iLID binding to the membrane of the synthetic cells was verified by imaging of the synthetic cells after addition of CTZ (Fig. 5b). Bioluminescence emission localized to the synthetic cells’ membrane was evident when imaging without laser excitation, and validated the binding and activity of Gluc (supplementary movie 1).

**Figure 5.**
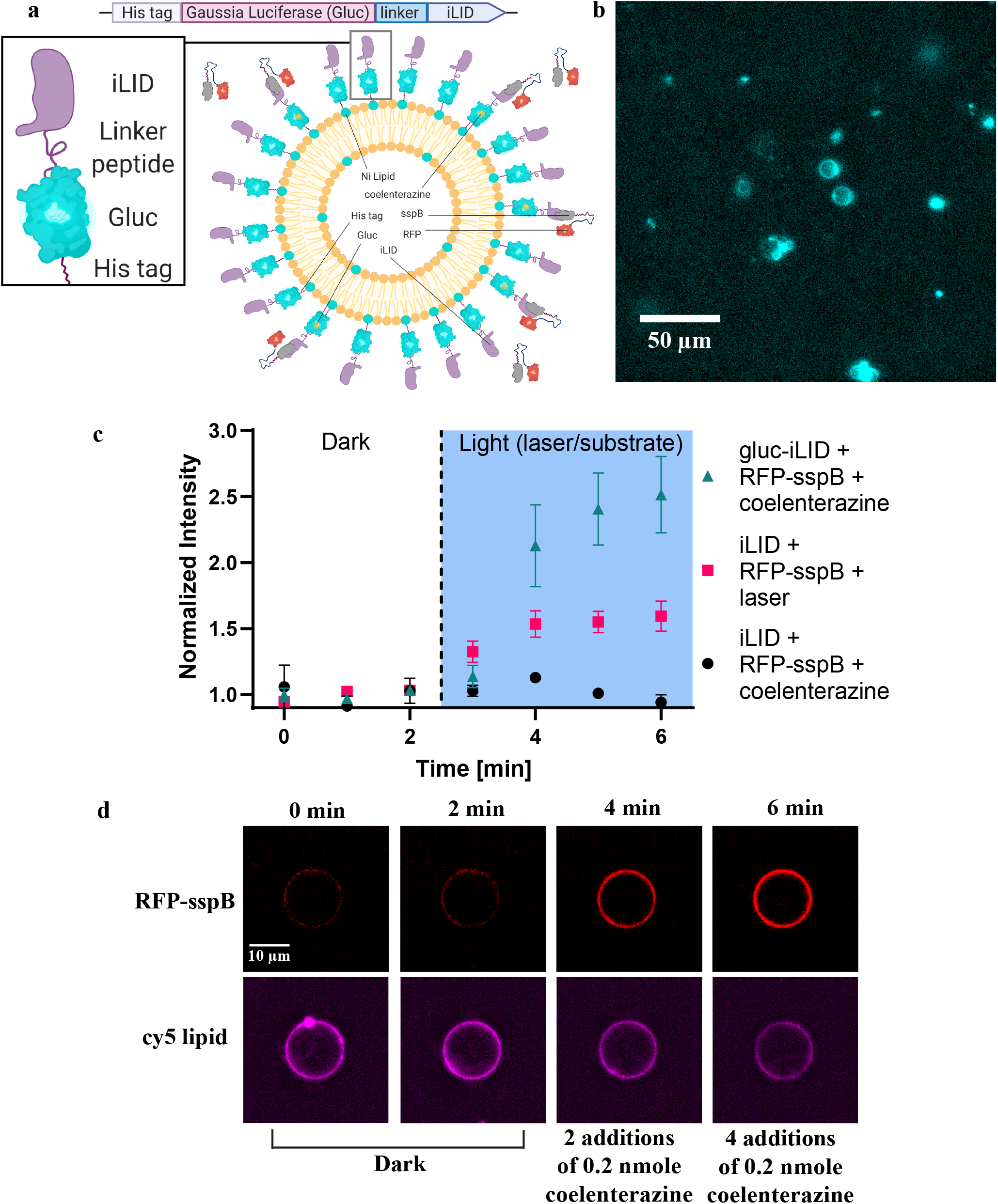
Bioluminescence-activated membrane recruitment in synthetic cells. **a,** A block diagram of the Gaussia luciferase (Gluc)-iLID fusion protein elements. Below, a schematic representation of protein recruitment to the synthetic cell membrane by hetero-dimerization of the fusion protein Gluc-iLID with RFP-sspB Nano. **b,** Microscopy image of luciferase light emission from synthetic cells with membrane-bound Gluc-EL222 after coelenterazine addition. **c,** Membrane recruitment of RFP-sspB Nano with iLID or Gluc-iLID using 488 nm laser illumination or by addition of coelenterazine to activate the bioluminescent reaction. RFP intensity is normalized to the average intensity of each cell in the dark conditions. Data is expressed as a mean ± s.e.m. (n=3 for iLID + coelenterazine, n=10 for iLID + laser, n=14 for Gluc-iLID + coelenterazine). **d,** Single-cell images of RFP-sspB Nano recruitment to a synthetic cell with membrane-bound Gluc-iLID in the dark and after two and four doses of 0.2 nmol coelenterazine (Top row). Bottom row displays the fluorescence of the synthetic cell’s DOPE-Cy5 lipid that composed 0.5 mol% of its membrane.

Next, the activation of membrane recruitment of mRFP1-sspB-Nano protein to individual synthetic cells was quantified using fluorescent microscopy. For the purpose of following individual cells before and after substrate addition, we immobilized the synthetic cells on the slide by adding a biotinylated lipid (DOPE-PEG4-biotin) to the synthetic cell membrane and coating the microscope slides with streptavidin. Synthetic cells remained bound to the surface after multiple additions and mixing of substrate. Activation of iLID and Gluc-iLID was tested using either 488 nm laser or by addition of CTZ. The normalized mRFP1 intensity in synthetic cells with membrane-bound Gluc-iLID increased by 2.5-times after 4 additions of CTZ in comparison to the initial dark conditions (Fig. 5c, green triangles, supplementary movie 2). Synthetic cells with membrane-bound iLID (without Gluc) were not activated by addition of CTZ (Fig. 5c, black circles), but demonstrated a 1.5-times increase in mRFP1 signal after 4 minutes of blue laser activation (Fig. 5c, pink squares).

The difference in the activation levels between iLID and Gluc-iLID synthetic cells can be associated with the addition of Gluc to the N-terminal side, that placed the iLID protein further away from the membrane and increased its exposure to the external solution. It was previously shown that binding of iLID to a membrane reduced its sspB-Nano binding ability in comparison to unbound iLID due to steric hindrance^41^. Hence, the addition of Gluc, which also acts as a membrane-distancing anchor, improved this issue. This was further validated by activation of Gluc-iLID with blue laser light, which resulted in an even higher increase in mRFP1 intensity, with more than 8-times difference between light and dark conditions, compared to a 1.5-times light-to-dark difference in the original iLID (Fig. S14). Taken together, these results demonstrate efficient and controlled bioluminescence-activated membrane recruitment of sspB-tagged proteins to Gluc-iLID labeled synthetic cells.

## Discussion

Developing an array of signaling and communication pathways is of high importance in synthetic cells to improve their integration with live cells and tissues, and build complex control circuits^42,43^. We utilized bioluminescent signaling for self-activation of synthetic cells and for communication between synthetic and natural cells. By optimizing the membrane composition and expression of Gluc in synthetic cells, we produced sufficient levels of time-integrated intensity to activate photoconidiation in an adjacent population of fungal *T. atroviride* cells. Moreover, to overcome light intensity limitations, novel transmitter-responder fusion proteins were designed to autoactivate transcription and membrane recruitment in synthetic cells using BRET, facilitating the activation of responder proteins with high light intensity demands.

Light is a powerful tool for controlling cellular processes in natural and synthetic cells alike due to the growing availability of genetically encodable optogenetic tools. For *in-vivo* applications, tissue attenuation of blue light either necessitates using far-red wavelengths or intrusive insertion of light sources that can deliver light close to the target site. While engineered red-shifted light-responding proteins or upconversion nanoparticles can be used to avoid using blue-light illumination, having a palette of spectral options is highly advantageous for developing sophisticated and multifunctional synthetic cells^44–46^. Moreover, these methods suffer from limitations such as restricted spectral shifting and possible changes in activity in the case of red-shifted proteins, tissue overheating and biodistribution challenges when using upconversion nanoparticles.

Here, we have shown that the light generated by Gluc-expressing synthetic cells is sufficient to activate sensitive proteins such as retinal rhodopsins (as the generated light can be seen with the naked eye) and biochemical pathways that are able to integrate photons over time, as in the case of *T. atroviride*. This is the first time, as far as we know, that bioluminescence was used for intercelullar communication between synthetic and natural cells. Expanding the use of this mechanism to different cell types, including eukaryotic cells with natural or engineered light-responsive elements, holds promise as a specific and tunable tool to control cell activities.

The phospholipid membrane plays an important role in the light interactions of the synthetic cell. This was especially evident in the UVB and UVC range, where we demonstrated the phospholipid membrane’s capability to protect DNA from radiation induced-damage. From an evolutionary aspect, these results might hint that lipids possibly had an evolutionary role as UV protecting agents in the prebiotic world which lacked an ozone layer and had high radiation levels^47^. Moreover, the lipid composition was shown to modify the light transmission through the membrane. Interestingly, the lipids’ hydrocarbon tail length and saturation level were previously shown to effect the membrane’s permeability to small molecules in a similar manner to that observed for light penetration in this study^48^. Therefore, light intensities going in and out of the synthetic cell can be tuned by selecting the appropriate membrane components.

Synthetic cells emit diffused light, in contrast to the focused light of lasers and LEDs, that are used traditionally to activate light-sensitive proteins in lab settings. Thus, close proximity between the synthetic cell and activated cell is required to compensate for this difference and achieve high local light intensity. Nonetheless, further improvements in the light-generating and light-responding elements are necessary to broaden the range of viable targets^49^. Possible avenues to increase light intensity include improving the enzyme and substrate performance, further optimizing the synthetic cell membrane for light transmission, and amplifying or focusing the generated signal. Moreover, engineering of the light-responsive proteins to increase their photosensitivity will contribute on the receiving end to enable this synthetic-natural cell cross-talk^50^.

Notably, using BRET enabled a stronger light response in synthetic cells by efficiently harnessing the energy generated by the bioluminescent reaction. These self-activating proteins can also be used for simultaneous imaging during the activation process, providing essential information on their activity site. The BRET approach expanded the range of light-responsive targets and facilitated the design and implementation of a bioluminescence-controlled transcription and membrane-recruitment mechanisms in synthetic cells. The controlled transcription mechanism for instance, could provide a possible tool for achieving precise temporal and spatial resolution of therapeutic protein expression *in-vivo*. While in this system, CTZ was supplied externally for initiation of the reaction, further exploration and incorporation of its biosynthesis will enable the next generation of synthetic cells to contain the complete signaling apparatus, and be used as independent light sources with therapeutic and diagnostic capabilities within the human body.

## Methods

### Lipids

1.2-dioleoyl-sn-glycero-3-phosphocholine (DOPC), 1-palmitoyl-2-oleoyl-sn-glycero-3-phosphocholine (POPC), 1,2-dimyristoyl-sn-glycero-3-phosphocholine (DMPC), 1.2-dipalmitoyl-sn-glycero-3-phosphocholine (DPPC), L-α-phosphatidylcholine, hydrogenated (Soy) (HSPC), 1,2-dioleoyl-sn-glycero-3-phosphoethanolamine (DOPE) and 1,2-dioleoyl-sn-glycero-3-phospho-(1’-rac-glycerol) (sodium salt) (DOPG) were purchased from Lipoid (Germany). 1,2-dioleoyl-sn-glycero-3-[(N-(5-amino-1-carboxypentyl) iminodiacetic acid) succinyl] (nickel salt) (DGS-NTA(Ni)) was purchased from Avanti Polar Lipids (USA). Cholesterol was purchased from Sigma-Aldrich (Israel). DOPE-PEG4-biotin and DOPE-cy5 were synthesized by reacting DOPE with NHS-PEG4-biotin or NHS-cy5 (BDL pharma, China).

### Synthetic cell preparation

#### Preparation of bacterial lysate for synthetic cell solutions

S30 bacterial lysate was prepared as described previously from BL21(DE3) *E. coli* transformed with the T7 polymerase expressing TargeTron vector pAR1219^51^. For Gaussia luciferase expressing synthetic cells, DTT and β-mercaptoethanol were excluded from the S30 solution during the lysate preparation.

#### Emulsion transfer method

Synthetic cell preparation using the emulsion transfer method was performed as previously described with several modifications^51^. The synthetic cells’ inner solution was either prepared according to the components lists in supplementary tables 3 and 4 or using the *E. coli* S30 extract system for circular DNA (Promega).

#### Shaking method

Synthetic cell preparation using the shaking method was performed as previously described by Gopfrich, et al^40^. 6mM of 100 nm liposomes were prepared using the thin film hydration method. A thin lipid film of POPC, DOPG, DGS-NTA(Ni), DOPE-PEG4-biotin and DOPE-Cy5 at molar ratios of 73.5:20:5:1:0.5 was hydrated with PBS containing 10 mM MgCl2 and extruded as described in the liposomes’ preparation section. 100 μl of liposomes diluted to 2 mM were mixed with 200 μl of FC-40 oil (FL-0005-HP; Iolitec, Germany) with 10 mM of PFPE–carboxylic acid (Krytox; Costenoble, Germany) and 0.8% of fluorosurfactant (RAN Biotechnologies, USA) and incubated overnight. The bottom layer was subsequently removed and 100 μl of PBS, followed by 100 μl of 1H,1H,2H,2H-Perfluoro-1-octanol (PFO; Sigma-Aldrich) were added on top of the remaining top layer. After 40 minutes of incubation, the top layer was extracted, containing the synthetic cells.

### UV exposure induced DNA damage

100 μg ml^−1^ of DNA (full sequence available in supplementary table 1) in PBS or DNA encapsulated in synthetic cells were exposed for 20 minutes to 254 nm UV light (R-52 Grid lamp; UVP, USA) placed 10 cm above a 96-well plate. The exposed free DNA was subsequently encapsulated in synthetic cells. DNA from the samples exposed to UV before and after encapsulation was extracted and PCR amplified using Phusion polymerase (Thermo-fisher, USA). The reaction products were then blotted on a 1% agarose gel.

### Luminescence assays

Luminescence was measured using the Infinite 200PRO multimode reader (TECAN). The CFPS or synthetic cells reactions were mixed in a 1:1 ratio with native coelenterazine (Nanolight, USA) just prior to the measurement.

#### Luciferase comparisons

For comparison of light production in Rluc and Gluc CFPS reactions with the unmodified internal (reducing) conditions, reactions were incubated at 37 °C and 1200 rpm for 1 hour. The reactions were measured after addition of a final concentration of 5 μM CTZ in a 384-well white microplate. For comparison of light production in Rluc synthetic cells and Gluc synthetic with the modified internal (oxidizing) conditions, synthetic cells were incubated at 37 °C for 1 hour. 40-fold diluted synthetic cells and CTZ in a final concentration of 5 μM were mixed and measured in a 384-well white microplate.

#### Luciferase Production kinetics

Gluc expressing synthetic cells were incubated at 37 °C. At each time point, 15 μl of synthetic cells were taken and diluted 4-fold in PBS. The diluted sample was measured after addition of a final concentration of 2.5 μM of CTZ in a 384-well white microplate.

#### Luciferase kinetics and re-activation of light emission

Gluc-expressing synthetic cells were incubated at 37 °C for 1 hour. For measuring the luciferase reaction kinetics, the synthetic cells were subsequently diluted 400-fold, mixed with a final concentration of 100 μM CTZ and measured in a 96-well white microplate. Luminescence was measured every 30 seconds for 15 minutes. For luminescence re-activation measurements, the synthetic cells were diluted 4-fold and mixed with 0.125 nmol of native CTZ in a 384-well white microplate. After measuring luminescence in one-minute intervals for 5 minutes, a second dose of 0.125 nmol of native CTZ was added to the same well and luminescence was measured again for 5 minutes.

### Synthetic cell size and concentration measurements

Size analysis of synthetic cells was performed by light diffraction using the Mastersizer 3000 (Malvern Instrument, UK). The synthetic cell concentration was measured by flow cytometry using the AMNIS ImageStream®^X^ Mk II (Luminex Corporation, USA). Synthetic cells were diluted 10-fold in the synthetic cells’ outer solution (supplementary table 5) and counted by dividing the number of events verified as synthetic cells using the images in the brightfield channel and the total volume analyzed by the device.

### Cryogenic scanning electron microscopy imaging

Cryogenic scanning electron microscopy (cryo-SEM) imaging we performed using Zeiss Ultra Plus high-resolution SEM, equipped with a Schottky field-emission gun and with a BalTec VCT100 cold-stage maintained below −145 °C. Specimens were imaged at low acceleration voltage of 1kV, and working distances of 3–5 mm. Both the Everhart Thornley (“SE2”) and the in-the-column (“InLens”) secondary electron imaging detectors were used. The energy-selective backscattered (“ESB”) detector was used for elemental contrast between the organic and the aqueous phases. Low-dose imaging was applied to all specimens to minimize radiation damages.

Specimens were prepared by the drop plunging method, a 3 μL drop of solution was set on top of a special planchette maintaining its droplet shape and was manually plunged into liquid ethane, after which it was set on top of a specialized sample table. The frozen droplets were transferred into the BAF060 freeze fracture system, where they were fractured by a rapid stroke from a cooled knife, exposing the inner part of the drop. They were then transferred into the pre-cooled HR-SEM as described above. Ideally, imaging was performed as close as possible to the drop surface, where cooling rate should be maximal.

### Protein expression and purification

BL21(DE3) *E. coli* (NEB) were used for the expression of DsbC-his, EL222-his, iLID and his-maltose binding protein (MBP)-mRFP1-sspB-Nano. Expression of Gluc-EL222-his and his-Gluc-iLID was performed in SHuffle T7 *E. coli*. A 5 ml Luria-broth starter culture for each protein was incubated overnight at 37 °C and 250 rpm with the compatible antibiotics (ampicillin at 100 μg ml^−1^ or kanamycin at 25 μg ml^−1^). The starter was then transferred to 500 ml of Terrific-broth supplemented with antibiotics in the same concentration and grown at 37 °C and 250 rpm to optical density (OD) of 0.5, when they were induced with 500 μM of Isopropyl β-D-1-thiogalactopyranoside (IPTG). DsbC-his and EL222-his were incubated at 37 °C and 250 rpm following induction until reaching an OD of ~4. His-iLID, his-MBP-mRFP1-sspB-Nano, Gluc-EL222-his and his-Gluc-iLID were grown at 16 °C and 250 rpm until reaching similar OD values. Cells were harvested by centrifugation at 7,000 × g for 10 minutes at 4 °C and kept at −20 °C until to the next step.

For protein purification, the pellet was resuspended in PBS (in the case of DsbC-his and EL222-his), 50 mM Tris, 300 mM NaCl, pH 7.4 (in the case of his-iLID and his-MBP-RFP-sspB-Nano) or 300 mM NaCl, 50 mM phosphate buffer, pH 8.0 (in the case of his-Gluc-iLID and Gluc-EL222-his). The cells were fractionated by two passes through an emulsiFlex-C3 high pressure homogenizer (Avestin, Germany) and the lysate was spun down two times for 15 minutes at 20,000 × g. The supernatant was passed through an AKTA purifier chromatography system (Cytiva, USA) using a HisTrap HP 5 ml column and eluted with elution buffer with similar composition to the loading buffer supplemented with 500 mM imidazole. The protein containing fractions were dialyzed in a 12-14 kD membrane (Spectrum Laboratories, USA) against their original resuspension buffer.

To remove the his-MBP domain from the mRFP1-SspB-Nano protein, the eluted MBP-RFP-sspB proteins were cut with TEV protease (NEB) using a digestion site between the MBP and the RPF sections. 300 μg of his-MBP-RFP-sspB-Nano were diluted to a total reaction volume of 880 μl. 20 μl of TEV Protease Reaction Buffer (10X) and 100 μl of TEV Protease were added, then incubated at 4°C overnight. 10 reactions were samples were pooled together and passed through a Ni Sepharose 6 Fast Flow histidine-tagged protein purification resin (Cytiva). The flow-through containing the RFP-SspB-Nano protein, was collected and concentrated using Amicon ultra 15 kDa (Merck, USA). The proteins were dialyzed overnight in PBS.

### Activation of photoconidiation in fungi with synthetic cells

#### Fungal growth

A *Trichoderma atroviride* inoculum, was plated on the center of a PDYC (24 g l^−1^ potato dextrose broth (Difco, UK), 1.2 g l^−1^ casein hydrolysate (Sigma-Aldrich), 2 g l^−1^ yeast extract (Difco) agar plate and incubated for twenty-four hours in dark conditions. 3 ml of PDYC media were subsequently added to a new 10 cm culture plate. In the center of the plate a Whatman 50 filter paper cut to a diameter of 9 cm was placed over an 8 cm Whatman 1 filter paper. On top of these, a 0.5 cm square from the periphery of the fungal growth on the agar plate was placed in the center and incubated in the dark at room temperature for 48 hours.

#### Exposure to synthetic cells

Synthetic cells were produced using the emulsion transfer method and diluted to the desired cell concentration. The synthetic cell concentration was quantified using flow cytometry using the AMNIS ImageStream®X Mk II as described above. Two consecutive exposures of the fungi to synthetic cells were performed. In each exposure, 500 μl of synthetic cells were mixed with 500 μl of native CTZ in a final concentration of 100 μM in a transparent plastic bag. The bag was localized over the periphery of the fungal colony on the top part of the plate for 15 minutes and then removed. The plates were left for incubation in the dark for an additional period 24 hours, after which they were imaged using a regular camera.

#### Image analysis

Background normalization of the images was performed manually in ImageJ to achieve the same average pixel value for all images^52^. The images were then converted to grayscale and then to black and white using a pixel threshold value equal to 45. The percentage of black pixels in a rectangle containing the top part of the plate where the synthetic cells were placed was calculated separately for each image.

### Transcription activation in synthetic cells

#### LED illumination system

Five 470 nm blue LEDs (C503B-BAN-CY0C0461; Mouser, Israel) were connected together in series using an Arduino microcontroller and an external power supplier 0-30 V. On-off intervals were set by controlling relay modules using the Arduino IDE software. Blue light intensity was measured with a S310C light sensor (Thorlabs, USA).

#### EL222 mediated activation of transcription in cell free reactions

Cell-free reactions based on the internal synthetic cell solution (supplementary table 3) for Rluc expression or *E. coli* S30 extract system for circular DNA (Promega) for RFP expression were supplemented with 2.5 μM EL222 (unless stated otherwise) and incubated at 37 °C in 384-well microplates coated with an adhesive film to prevent evaporation. LEDs were placed 7 cm above the plate, providing approximately 12 W m^−2^, with on-off intervals of 20 and 70 seconds. For analysis of Rluc expression, native CTZ was added to a final concentration of 5 μM and luminescence was measured. For analysis of RFP expression, fluorescence intensity was measured with excitation/emission at 540 nm / 600 nm.

#### EL222 activation in Synthetic cells

Synthetic cells with an internal solution based on the *E. coli* S30 extract system for circular DNA (Promega) were supplemented with 2.5 μM EL222. Prior to their incubation, each synthetic cell batch was divided into dark and light groups, both incubated at 37 °C in 384-well microplates coated with an adhesive film under dark or blue-light conditions. LEDs were placed 2.8 cm above the plate, providing approximately 19 W m^−2^, with on-off intervals of 20 and 70 seconds.

Self-activating synthetic cells based on the *E. coli* S30 extract system for circular DNA (Promega) were supplemented with 2.5 μM of Gluc-EL222. Prior to their incubation, each synthetic cell batch was divided into dark and light groups, both incubated at 37 ° C. Native CTZ (0.2 nmol) was added every 30 minutes to the light group for a total of four times. The samples were further incubated for 90 more minutes after the last substrate addition. Just prior to the final fluorescence measurements, CTZ was added to the dark group in the same concentration that was added to the light group in order to eliminate differences due to substrate auto-fluorescence.

### Membrane recruitment in synthetic cells

#### Membrane recruitment of RFP-sspB-Nano

A PDMS-walled chamber (Sylgard 184; Dow, USA), 5.5 mm in diameter and 2.5 mm height, was placed on a 22 × 50 mm deckglaser glass slides (Marienfeld, Germany). The bottom of the chamber was coated with 10 μg ml^−1^ streptavidin (Promega) overnight at 4°C, or with 1% bovine serum albumin (Sigma-Aldrich) for 1 hour at room temperature. The chamber was washed with PBS and in case of streptavidin coating, was coated once more for 1 hour with 5 mg ml^−1^ of cold water fish skin gelatin (Sigma-Aldrich) and washed again with PBS.

Synthetic cells were prepared using the shaking method (see synthetic-cell preparation), and 100 nM of his-iLID or his-Gluc-iLID were added to the solution and incubated for 1 hour at room temperature and shaking of 100 rpm. Under dark conditions, 25 nM of RFP-sspB-Nano were added to the cells and placed in the coated chamber in dark conditions for 30 minutes.

Imaging was performed using a standard inverted microscope (Eclipse Ti2; Nikon) outfitted with a 60x 1.4 NA objective lens (Nikon). Low power laser illumination of approximately 5 mW cm^−2^ 640 nm and 500 mW cm^−2^ 561 nm lasers for imaging of Cy5 and mRFP1 respectively were used. Images were captured on a Sona sCMOS detector (Andor, Northern Ireland). Blue light laser illumination was performed with 488 nm laser at approximately 20 mW cm^−2^ with on-off intervals of 1.25 and 3.75 seconds for one minute. For recruitment in synthetic cells functionalized with his-Gluc-iLID, native coelenterazine (0.2 nmol) was added every minute for a total of four minutes.

#### Gluc-iLID light emission imaging

Synthetic cells functionalized with Gluc-iLID were imaged after addition of 100 μM substrate (final concentration) using a standard inverted microscope (Eclipse Ti2; Nikon) outfitted with a 40x 0.75 NA objective lens (Nikon) and equipped with iXON EMCCD camera (Andor). Images were obtained with 400 msec exposure time and gain 300.

### Statistics

The statistical analysis including student’s t-test, one-way and two-way ANOVA was performed using Prism GraphPad version 8.3.

## Acknowledgments

This project received funding from the European Union’s Horizon 2020 research and innovation program under the grant agreement No 680242-ERC-[Next-Generation Personalized Diagnostic Nanotechnologies for Predicting Response to Cancer Medicine]. The authors also acknowledge the support of Israel Innovation Authority for the Nofar Grant (67967), the Israel Science Foundation (1778/13, 1421/17); The Israel Ministry of Economy for a Kamin Grant (52752, 69230); the Israel Ministry of Science, Technology & Space (3-16963; 3-17418); the Ministry of Agriculture & Rural Development - Office of the Chief Scientist (323/19); the Israel Cancer Association (2015-0116); the Leventhal 2020 COVID19 Research Fund (ATS #11947), the German-Israeli Foundation for Scientific Research and Development for a GIF Young grant (I-2328-1139.10/2012); the European Union FP-7 IRG Program for a Career Integration Grant (908049); the Phospholipid Research Center Grant (ASC-2018-062/1-1); the European Research Council (ERC) under the European Union Horizon 2020 research and innovation programme to Y.S. (802567); the European Research Council (ERC) to L.G. (ERC-2017-COG-773181-iPS-ChOp-AF); the Louis family Cancer Research Fund, a Mallat Family Foundation Grant; the Unger Family Fund; a Carrie Rosenblatt Cancer Research Fund, the Technion Integrated Cancer Center (TICC), the Russell Berrie Nanotechnology Institute, and the Lorry I. Lokey Interdisciplinary Center for Life Sciences & Engineering. A.S. acknowledges the Alon and Taub Fellowships. O.A. acknowledges the Jacobs and Gutwirth Fellowships. L.E.W. and Y.S. acknowledges the Zuckerman foundation, G.C. acknowledges the “Baroness Ariane de Rothschild Women Doctoral Program” from the Rothschild Caesarea Foundation. The Max Planck Society is appreciated for its general support.

The authors thank Mrs. Naama Koifman from the Technion Center for Electron Microscopy of Soft Matter (TCEMSM) for performing the cryo-SEM imaging, and Mr. Nadav Opatovski for his technical assistance with microscope calibration. The authors thank Dr. Assaf Zinger (Department of Chemical Engineering, Technion), Dr. Shai Berlin, Mr. Michael Andreyanov (Department of Medicine, Technion), Prof. Ygal Rotenstreich, Dr. Ifat Sher, Mrs. Zehavit Goldberg (Sheba Medical Center, Israel) for their professional input and helpful discussions that greatly contributed to the paper. Illustrations in this paper were created with BioRender.com and Adobe Illustrator.

