## supplementary material for "Bioluminescent Synthetic Cells Communicate with Natural Cells and Self-Activate Light-Responsive Proteins"

### Supplementary Table of Contents

### Supplementary Notes

#### Plasmids and hosts

The DNA sequences of all the proteins used in this study are listed in supplementary table 1. *E. coli* DH5α and TOP10 strains were used for cloning and plasmid purification. DNA sequences of the engineered plasmids were confirmed by sequencing. A plasmid expressing *Renilla* luciferase (Rluc) under the T7 promoter was obtained from the S30-T7 high yield protein expression system kit, purchased from Promega (USA). *Gaussia* luciferase (Gluc) expressing plasmid was generously provided Prof. James Swartz (department of chemical engineering, Stanford university). Plasmids expressing DsbC (Plasmid #38152), EL222 (Plasmid #113108), iLID (Plasmid #60408), sspB-Nano (Plasmid #60409), and pBLind RFP (Plasmid #113109) were purchased from Addgene (USA). PCR was used to add a C-terminal his tag to the DsbC protein and insert it to a pET28a vector. C-terminal his tag was also added to the EL222 sequence that was isolated and inserted into a pET28a vector. Hifi DNA assembly (NEB, USA) was used to produce Gluc-EL222-his and his-Gluc-iLID sequences, each inserted to a pet28a vector. mRFP1 sequence was inserted between the MBP and sspB-Nano reading frames to generate a his-MBP-RFP-sspB-Nano vector. A plasmid expressing Rluc under the pBLind promoter was produced by replacing the RFP sequence in the original vector with the Rluc sequence from the Rluc expressing plasmid. A complete list of the primers used for plasmid design is available in supplementary table 2.

#### Preparation of lipids in mineral oil

POPC was lyophilized (FreeZone 2.5; Labconco, USA) overnight. POPC and cholesterol were dissolved separately in chloroform at a concentration of 50 mg ml$^{-1}$ each. 50 µl of each solution was added to 500 µl mineral oil (Sigma-Aldrich) in a glass vial. The mixture was vortexed and then heated at 80 °C for one hour. The obtained lipids oil was stored at room temperature for up to 2 weeks.

#### Liposome preparation and absorbance measurements

Liposomes were prepared using the ethanol injection method. Lipids were weighed and dissolved in absolute ethanol and subsequently injected into calcium-free Dulbecco's phosphate buffer saline (PBS; Sigma-Aldrich) preheated to 65 ºC reaching a final lipid concentration of 50 mM. The liposomes were extruded five times using a high-pressure Lipex extruder (Northern Lipids, Canada) through 400, 200, and 100 nm polycarbonate-etched membrane (Whatman, Newton, MA, USA) at 65 ºC.

For the absorbance measurements, liposomes were diluted 50-fold and measured in a UV-star 96-well microplate (Greiner bio-one, Germany) using the Infinite 200PRO multimode reader (TECAN, Switzerland).

**Western blot Analysis**

Synthetic cell samples after protein production were diluted 8-fold and analyzed using SDS-PAGE with a 12% acrylamide gel. The gel was blotted onto a nitrocellulose membrane and blocked with 5% nonfat milk powder in Tris-buffered saline. *Gaussia* luciferase Polyclonal Antibody (Invitrogen, USA) diluted 1:3750 in Tris-buffered saline with 0.5% Tween-20 and 0.5% nonfat milk powder was incubated with the membrane overnight at 4 °C. After washing, the blots were incubated with horseradish peroxidase-conjugated anti-rabbit (goat origin) secondary antibody (GenScript, USA) diluted to 1:20,000 and developed with Clarity Western ECL Blotting Substrate (BioRad, USA). The results were imaged using the Fusion FX6 imaging system (Vilber, France).

For quantification of Gluc production, analysis of the images was performed with ImageJ gel analysis plug-in and use of a calibration curve for Gluc in known concentrations.

### Supplementary Tables

| Name | Sequence |
|---|---|
| *Renilla* Luciferase | ATGGCTTCCAAGGTGTACGACCCCGAGCAACGCAAACGCATGATCACTGGGCCTCAGTGGTGG<br>GCTCGCTGCAAGCAAATGAACGTGCTGGACTCCTTCATCAACTACTATGATTCCGAGAAGCAC<br>GCCGAGAACGCCGTGATTTTTCTGCATGGTAACGCTGCCTCCAGCTACCTGTGGAGGCACGTC<br>GTGCCTCACATCGAGCCCGTGGCTAGATGCATCATCCCTGATCTGATCGGAATGGGTAAGTCC<br>GGCAAGAGCGGGAATGGCTCATATCGCCTCCTGGATCACTACAAGTACCTCACCGCTTGGTTC<br>GAGCTGCTGAACCTTCCAAAGAAAATCATCTTTGTGGGCCACGACTGGGGGGCTTGTCTGGCC<br>TTTCACTACTCCTACGAGCACCAAGACAAGATCAAGGCCATCGTCCATGCTGAGAGTGTCGTG<br>GACGTGATCGAGTCCTGGGACGAGTGGCCTGACATCGAGGAGGATATCGCCCTGATCAAGAGC<br>GAAGAGGGCGAGAAAATGGTGCTTGAGAATAACTTCTTCGTCGAGACCATGCTCCCAAGCAAG<br>ATCATGCGGAAACTGGAGCCTGAGGAGTTCGCTGCCTACCTGGAGCCATTCAAGGAGAAGGGC<br>GAGGTTAGACGGCCTACCCTCTCCTGGCCTCGCGAGATCCCTCTCGTTAAGGGAGGCAAGCCC |

| | |
|---|---|
| | GACGTCGTCCAGATTGTCCGCAACTACAACGCCTACCTTCGGGCCAGCGACGATCTGCCTAAG ATGTTCATCGAGTCCGACCCTGGGTTCTTTTCCAACGCTATTGTCGAGGGAGCTAAGAAGTTCC CTAACACCGAGTTCGTGAAGGTGAAGGGCCTCCACTTCAGCCAGGAGGACGCTCCAGATGAAA TGGGTAAGTACATCAAGAGCTTCGTGGAGCGCGTGCTGAAGAACGAGCAGTAA |
| *Gaussia* Luciferase | ATGAAACCAACAGAAAATAACGAAGATTTTAACATCGTCGCGGTAGCCTCAAATTTTGCAACC ACCGACTTAGACGCGGATCGTGGGAAATTACCAGGTAAGAAATTGCCGTTGGAAGTACTTAAA GAACTGGAAGCCAACGCACGCAAGGCTGGATGCACTCGTGGGTGTTTGATCTGTCTGTCGCAT ATTAAGTGTACGCCAAAGATGAAAAAGTTCATCCCAGGGCGTTGTCATACTTACGAAGGAGAC AAAGAGTCAGCACAGGGCGGAATCGGGGAAGCTATCGTAGATATCCCTGAGATCCCAGGCTTC AAAGACTTGGAACCGCTGGAACAGTTTATTGCTCAAGTTGACTTGTGCGTCGACTGTACGACT GGGTGCCTTAAAGGGCTGGCGAATGTCCAATGTAGTGATCTGCTTAAGAAGTGGCTTCCTCAG CGTTGCGCTACATTCGCAAGCAAAATTCAGGGGCAAGTGGATAAGATCAAAGGCGCGGGTGG TGACTAA |
| EL222-his | ATGGGTATGTTGGATATGGGACAAGATCGGCCGATCGATGGAAGTGGGGCACCCGGGGCAGA CGACACACGCGTTGAGGTGCAACCGCCGGCGCAGTGGGTCCTCGACCTGATCGAGGCCAGCCC GATCGCATCGGTCGTGTCCGATCCGCGTCTCGCCGACAATCCGCTGATCGCCATCAACCAGGC CTTCACCGACCTGACCGGCTATTCCGAAGAAGAATGCGTCGGCCGCAATTGCCGATTCCTGGC AGGTTCCGGCACCGAGCCGTGGCTGACCGACAAGATCCGCCAAGGCGTGCGCGAGCACAAGC CGGTGCTGGTCGAGATCCTGAACTACAAGAAGGACGGCACGCCGTTCCGCAATGCCGTGCTCG TTGCACCGATCTACGATGACGACGACGAGCTTCTCTATTTCCTCGGCAGCCAGGTCGAAGTCG ACGACGACCAGCCCAACATGGGCATGGCGCGCCGCGAACGCGCCGCGGAAATGCTCAAGACG CTGTCGCCGCGCCAGCTCGAGGTTACGACGCTGGTGGCATCGGGCTTGCGCAACAAGGAAGTG GCGGCCCGGCTCGGCCTGTCGGAGAAAACCGTCAAGATGCACCGCGGGCTGGTGATGGAAAA GCTCAACCTGAAGACCAGCGCCGATCTGGTGCGCATTGCCGTCGAAGCCGGAATCCATCATCA TCATCATCACTAA |
| his-iLID | ATGAGAGGATCGCATCACCATCACCATCACGGATCCGGGGAGTTTCTGGCAACCACACTGGAA CGGATCGAGAAAAATTTCGTGATTACTGATCCGAGACTGCCTGACAACCCAATCATTTTTGCG AGCGATTCCTTCCTGCAGCTGACAGAATATTCTCGGGAAGAGATCCTGGGGCGCAATTGCCGT TTTCTGCAGGGACCCGAGACAGACCGTGCCACTGTTCGGAAAATCAGAGATGCTATTGACAAC CAGACTGAAGTGACCGTTCAGCTGATCAATTATACCAAGAGCGGCAAGAAGTTCTGGAACGTG TTCCACCTGCAGCCGATGCGCGATTATAAGGGCGACGTCCAGTACTTCATTGGCGTGCAGCTG GATGGCACCGAACGTCTTCATGGCGCCGCTGAGCGTGAGGCGGTCATGCTGATCAAAAAGACA GCCTTTCAGATTGCTGAGGCAGCGAACGACGAAAATTACTTTTAA |
| his-MBP-RFP-sspB-Nano | ATGAGAGGATCGCATCACCATCACCATCACGGATCTAAAATCGAAGAAGGTAAACTGGTAATC TGGATTAACGGCGATAAAGGCTATAACGGTCTCGCTGAAGTCGGTAAGAAATTCGAGAAAGAT ACCGGAATTAAAGTCACCGTTGAGCATCCGGATAAACTGGAAGAGAAATTCCCACAGGTTGCG GCAACTGGCGATGGCCCTGACATTATCTTCTGGGCACACGACCGCTTTGGTGGCTACGCTCAAT CTGGCCTGTTGGCTGAAATCACCCCGGACAAAGCGTTCCAGGACAAGCTGTATCCGTTTACCT GGGATGCCGTACGTTACAACGGCAAGCTGATTGCTTACCCGATCGCTGTTGAAGCGTTATCGCT GATTTATAACAAAGACCTGCTGCCGAACCCGCCAAAAACCTGGGAAGAGATCCCGGCGCTGG ATAAAGAACTGAAAGCGAAAGGTAAGAGCGCGCTGATGTTCAACCTGCAAGAACCGTACTTC ACCTGGCCGCTGATTGCTGCTGACGGGGGTTATGCGTTCAAGTATGAAAACGGCAAGTACGAC ATTAAAGACGTGGGCGTGGATAACGCTGGCGCGAAAGCGGGTCTGACCTTCCTGGTTGACCTG ATTAAAAACAAACACATGAATGCAGACACCGATTACTCCATCGCAGAAGCTGCCTTTAATAAA GGCGAAACAGCGATGACCATCAACGGCCCGTGGGCATGGTCCAACATCGACACCAGCAAAGT GAATTATGGTGTAACGGTACTGCCGACCTTCAAGGGTCAACCATCCAAACCGTTCGTTGGCGT GCTGAGCGCAGGTATTAACGCCGCCAGTCCGAACAAAGAGCTGGCAAAAGAGTTCCTCGAAA ACTATCTGCTGACTGATGAAGGTCTGGAAGCGGTTAATAAAGACAAACCGCTGGGTGCCGTAG CGCTGAAGTCTTACGAGGAAGAGTTGGCGAAAGATCCACGTATTGCCGCCACTATGGAAAACG CCCAGAAAGGTGAAATCATGCCGAACATCCCGCAGATGTCCGCTTTCTGGTATGCCGTGCGTA CTGCGGTGATCAACGCCGCCAGCGGTCGTCAGACTGTCGATGAAGCCCTGAAAGACGCGCAGA CTAATTCGAGCTCGAACAACAACAACAATAACAATAACAACAACCTCGGGATCGAGGGAACG ACCGAAAACCTGTATTTTCAGGGATCCATGGCGAGTAGCGAAGACGTTATCAAAGAGTTCATG CGTTTCAAAGTTCGTATGGAAGGTTCCGTTAACGGTCACGAGTTCGAAATCGAAGGTGAAGGT |

| | |
|---|---|
| | GAAGGTCGTCCGTACGAAGGTACCCAGACCGCTAAACTGAAAGTTACCAAAGGTGGTCCGCTG<br>CCGTTCGCTTGGGACATCCTGTCCCCGCAGTTCCAGTACGGTTCCAAAGCTTACGTTAAACACC<br>CGGCTGACATCCCGGACTACCTGAAACTGTCCTTCCCGGAAGGTTTCAAATGGGAACGTGTTA<br>TGAACTTCGAAGACGGTGGTGTTGTTACCGTTACCCAGGACTCCTCCCTGCAAGACGGTGAGTT<br>CATCTACAAAGTTAAACTGCGTGGTACCAACTTCCCGTCCGACGGTCCGGTTATGCAGAAAAA<br>AACCATGGGTTGGGAAGCTTCCACCGAACGTATGTACCCGGAAGACGGTGCTCTGAAAGGTGA<br>AATCAAAATGCGTCTGAAACTGAAAGACGGTGGTCACTACGACGCTGAAGTTAAAACCACCTA<br>CATGGCTAAAAAACCGGTTCAGCTGCCGGGTGCTTACAAAACCGACATCAAACTGGACATCAC<br>CTCCCACAACGAAGACTACACCATCGTTGAACAGTACGAACGTGCTGAAGGTCGTCACTCCAC<br>CGGTGCTCTGCAGAGCTCCCCGAAACGCCCTAAGCTGCTGCGTGAATATTACGATTGGCTGGTT<br>GATAACAGCTTTACCCCATATCTGGTGGTGGATGCCACATACCTGGGCGTGAACGTGCCCGTG<br>GAGTATGTGAAAGACGGTCAGATCGTGCTGAATCTGTCTGCAAGTGCGACCGGCAACCTGCAA<br>CTGACAAATGATTTTATCCAGTTCAACGCCCGCTTTAAGGGCGTGTCTCGTGAACTGTATATCC<br>CGATGGGTGCCGCTCTGGCCATTTACGCTCGCGAGAACGGCGATGGTGTGATGTTCGAACCAG<br>AAGAAATCTATGACGAGCTGAATATTGGTTAA |
| DsbC-his | ATGGCTAGCATGACTGGTGGACAGCAAATGGGTCGGATCATCACAAGTTTGTACAAAAAAGTT<br>GTGGACGACGCGGCAATTCAACAAACGTTAGCCAAAATGGGCATCAAAAGCAGCGATATTCA<br>GCCCGCGCCTGTAGCTGGCATGAAGACAGTTCTGACTAACAGCGGCGTGTTGTACATCACCGA<br>TGATGGTAAACATATCATTCAGGGGCCAATGTATGACGTTAGTGGCACGGCTCCGGTCAATGT<br>CACCAATAAGATGCTGTTAAAGCAGTTGAATGCGCTTGAAAAAGAGATGATCGTTTATAAAGC<br>GCCGCAGGAAAAACACGTCATCACCGTGTTTACTGATATTACCTGTGGTTACTGCCACAAACT<br>GCATGAGCAAATGGCAGACTACAACGCGCTGGGGATCACCGTGCGTTATCTTGCTTTCCCGCG<br>CCAGGGGCTGGACAGCGATGCAGAGAAAGAAATGAAAGCTATCTGGTGTGCGAAAGATAAAA<br>ACAAAGCGTTTGATGATGTGATGGCAGGTAAAAGCGTCGCACCAGCCAGTTGCGACGTGGATA<br>TTGCCGACCATTACGCACTTGGCGTCCAGCTTGGCGTTAGCGGTACTCCGGCAGTTGTGCTGAG<br>CAATGGCACACTTGTTCCGGGTTACCAGCCGCCGAAAGAGATGAAAGAATTCCTCGACGAACA<br>CCAAAAAATGACCAGCGGTAAACACCACCACCACCACCACTAA |
| his-*Gaussia* Luciferase-iLID | ATGCATCACCATCACCATCACAAACCAACAGAAAATAACGAAGATTTTAACATCGTCGCGGTA<br>GCCTCAAATTTTGCAACCACCGACTTAGACGCGGATCGTGGGAAATTACCAGGTAAGAAATTG<br>CCGTTGGAAGTACTTAAAGAACTGGAAGCCAACGCACGCAAGGCTGGATGCACTCGTGGGTGT<br>TTGATCTGTCTGTCGCATATTAAGTGTACGCCAAAGATGAAAAAGTTCATCCCAGGGCGTTGTC<br>ATACTTACGAAGGAGACAAAGAGTCAGCACAGGGCGGAATCGGGGAAGCTATCGTAGATATC<br>CCTGAGATCCCAGGCTTCAAAGACTTGGAACCGCTGGAACAGTTTATTGCTCAAGTTGACTTGT<br>GCGTCGACTGTACGACTGGGTGCCTTAAAGGGCTGGCGAATGTCCAATGTAGTGATCTGCTTA<br>AGAAGTGGCTTCCTCAGCGTTGCGCTACATTCGCAAGCAAAATTCAGGGGCAAGTGGATAAGA<br>TCAAAGGCGCGGGTGGTGACGGTGGTGGTGGTTCAGGTGGTGGTGGTTCAGGATCCGGGGAGT<br>TTCTGGCAACCACACTGGAACGGATCGAGAAAAATTTCGTGATTACTGATCCGAGACTGCCTG<br>ACAACCCAATCATTTTTGCGAGCGATTCCTTCCTGCAGCTGACAGAATATTCTCGGGAAGAGAT<br>CCTGGGGCGCAATTGCCGTTTTCTGCAGGGACCCGAGACAGACCGTGCCACTGTTCGGAAAAT<br>CAGAGATGCTATTGACAACCAGACTGAAGTGACCGTTCAGCTGATCAATTATACCAAGAGCGG<br>CAAGAAGTTCTGGAACGTGTTCCACCTGCAGCCGATGCGCGATTATAAGGGCGACGTCCAGTA<br>CTTCATTGGCGTGCAGCTGGATGGCACCGAACGTCTTCATGGCGCCGCTGAGCGTGAGGCGGT<br>CATGCTGATCAAAAAGACAGCCTTTCAGATTGCTGAGGCAGCGAACGACGAAAATTACTTTTA<br>A |
| *Gaussia* Luciferase-EL222-his | ATGAAACCAACAGAAAATAACGAAGATTTTAACATCGTCGCGGTAGCCTCAAATTTTGCAACC<br>ACCGACTTAGACGCGGATCGTGGGAAATTACCAGGTAAGAAATTGCCGTTGGAAGTACTTAAA<br>GAACTGGAAGCCAACGCACGCAAGGCTGGATGCACTCGTGGGTGTTTGATCTGTCTGTCGCAT<br>ATTAAGTGTACGCCAAAGATGAAAAAGTTCATCCCAGGGCGTTGTCATACTTACGAAGGAGAC<br>AAAGAGTCAGCACAGGGCGGAATCGGGGAAGCTATCGTAGATATCCCTGAGATCCCAGGCTTC<br>AAAGACTTGGAACCGCTGGAACAGTTTATTGCTCAAGTTGACTTGTGCGTCGACTGTACGACT<br>GGGTGCCTTAAAGGGCTGGCGAATGTCCAATGTAGTGATCTGCTTAAGAAGTGGCTTCCTCAG<br>CGTTGCGCTACATTCGCAAGCAAAATTCAGGGGCAAGTGGATAAGATCAAAGGCGCGGGTGG<br>TGACGGTGGTGGTGGTTCAGGTGGTGGTGGTTCAATGTTGGATATGGGACAAGATCGGCCGAT<br>CGATGGAAGTGGGGCACCCGGGGCAGACGACACACGCGTTGAGGTGCAACCGCCGGCGCAGT |

| | |
|---|---|
| | GGGTCCTCGACCTGATCGAGGCCAGCCCGATCGCATCGGTCGTGTCCGATCCGCGTCTCGCCG ACAATCCGCTGATCGCCATCAACCAGGCCTTCACCGACCTGACCGGCTATTCCGAAGAAGAAT GCGTCGGCCGCAATTGCCGATTCCTGGCAGGTTCCGGCACCGAGCCGTGGCTGACCGACAAGA TCCGCCAAGGCGTGCGCGAGCACAAGCCGGTGCTGGTCGAGATCCTGAACTACAAGAAGGAC GGCACGCCGTTCCGCAATGCCGTGCTCGTTGCACCGATCTACGATGACGACGACGAGCTTCTCT ATTTCCTCGGCAGCCAGGTCGAAGTCGACGACGACCAGCCCAACATGGGCATGGCGCGCCGCG AACGCGCCGCGGAAATGCTCAAGACGCTGTCGCCGCGCCAGCTCGAGGTTACGACGCTGGTGG CATCGGGCTTGCGCAACAAGGAAGTGGCGGCCCGGCTCGGCCTGTCGGAGAAAACCGTCAAG ATGCACCGCGGGCTGGTGATGGAAAAGCTCAACCTGAAGACCAGCGCCGATCTGGTGCGCATT GCCGTCGAAGCCGGAATCCATCATCATCATCATCACTAA |
| mRFP1 | ATGGCGAGTAGCGAAGACGTTATCAAAGAGTTCATGCGTTTCAAAGTTCGTATGGAAGGTTCC GTTAACGGTCACGAGTTCGAAATCGAAGGTGAAGGTGAAGGTCGTCCGTACGAAGGTACCCA GACCGCTAAACTGAAAGTTACCAAAGGTGGTCCGCTGCCGTTCGCTTGGGACATCCTGTCCCC GCAGTTCCAGTACGGTTCCAAAGCTTACGTTAAACACCCGGCTGACATCCCGGACTACCTGAA ACTGTCCTTCCCGGAAGGTTTCAAATGGGAACGTGTTATGAACTTCGAAGACGGTGGTGTTGTT ACCGTTACCCAGGACTCCTCCCTGCAAGACGGTGAGTTCATCTACAAAGTTAAACTGCGTGGT ACCAACTTCCCGTCCGACGGTCCGGTTATGCAGAAAAAAACCATGGGTTGGGAAGCTTCCACC GAACGTATGTACCCGGAAGACGGTGCTCTGAAAGGTGAAATCAAAATGCGTCTGAAACTGAA AGACGGTGGTCACTACGACGCTGAAGTTAAAACCACCTACATGGCTAAAAAACCGGTTCAGCT GCCGGGTGCTTACAAAACCGACATCAAACTGGACATCACCTCCCACAACGAAGACTACACCAT CGTTGAACAGTACGAACGTGCTGAAGGTCGTCACTCCACCGGTGCTTAA |
| pBLind promoter | GGTAGCCTTTAGTCCATGTTAGCGAAGAAAATGGTTTGTTATAGTCGAATAAA |
| UV exposed oligo | GGCGCAATCACGAATGAATAACGGTTTGGTTGATGCGAGTGATTTTGATGACGAGCGTAATGG CTGGCCTGTTGAACAAGTCTGGAAAGAAATGCATAAACTTTTGCCATTCTCACCGGATTCAGTC GTCACTCATGGTGATTTCTCACTTGATAACCTTATTTTTGACGAGGGGAAATTAATAGGTTGTA TTGATGTTGGACGAGTCGGAATCGCAGACCGATACCAGGATCTTGCCATCCTATGGAACTGCC TCGGTGAGTTTTCTCCTTCATTACAGAAACGGCTTTTTCAAAAATATGGTATTGATAATCCTGA TATGAATAAATTGCAGTTTCATTTGATGCTCGATGAGTTTTTCTAAGAATTAATTCATGAGCGG ATACATATTTGAATGTATTTAGAAAAATAAACAAATAGGGGTTCCGCGCACATTTCCCCGAAA AGTGCCACCTGAAATTGTAAACGTTAATATTTTGTTAAAATTCGCGTTAAATTTTTGTTAAATC AGCTCATTTTTTAACCAATAGGCCGAAATCGGCAAAATCCCTTATAAATCAAAAGAATAGACC GAGATAGGGTTGAG |

**Supplementary Table 1.** Protein and promoter sequences.

| **Primer** | **Sequence** |
|---|---|
| DsbC forward primer | TAATACGACTCACTATAGGG |
| DsbC reverse primer | GGATCCTTAGTGGTGGTGGTGGTGGTGTTTACCGCTGGTCAT |
| EL222 forward primer | ATGCGCCATGGGTATGTTGGATATGGGACAAGATCGG |
| EL222 reverse primer | ATCGCGGATCCTTAGTGATGATGATGATGATGGATTCCGGCTTCGACGG CAA |
| mRFP1 forward primer | CAGTCGGATCCATGGCGAGTAGCGAAGACGT |
| mRFP1 reverse primer | AGCATCTGCAGAGCACCGGTGGAGTGACGA |
| pQE-sspB forward primer | AATGACTGCAGAGCTCCCCGAAACGCCCTAA |
| pQE-sspB reverse primer | GGAGCTGGATCCCTGAAAATACAGG |
| UV oligo forward primer | GGCGCAATCACGAATGAATA |
| UV oligo reverse primer | CTCAACCCTATCTCGGTCTA |
| iLID forward primer | GGATCCGGGGAGTTTCTGGC |
| iLID reverse primer | CAAGCTTGTCGACGGAGCTCGAATTCTTAAAAGTAATTTTCGTCGTTCGC TGC |
| *Gaussia* luciferase forward primer | GTTTAACTTTAAGAAGGAGATATACATATGCATCACCATCACCATCACA AAC |

| | |
|---|---|
| *Gaussia* luciferase reverse primer | GTCACCACCCGCGCCTTTGATC |
| *Gaussia* luciferase-iLID-linker | TAAGATCAAAGGCGCGGGTGGTGACGGTGGTGGTGGTTCAGGTGGTGGT GGTTCAGGATCCGGGGAGTTTCTGGCAACCA |
| *Gaussia* luciferase-EL222 forward primer | TTTTGTTTAACTTTAAGAAGGAGATATACATATGAAACC |
| *Gaussia* luciferase-linker-rev | TCTTGTCCCATATCCAACATTGAACCACCACCACCTGAACCACCACCAC CGTCACCACCCGCGCCTTT |
| *Renilla* luciferase forward primer | ATAGACATATGGCTTCCAAGGTGTACGA |
| *Renilla* luciferase reverse primer | GATTTGGATCCTTACTGCTCGTTCTTCAGCA |

**Supplementary Table 2.** Primers used for plasmid design.

| **Reagent** | **final concentration** | |
|---|---|---|
| Sucrose | 200 | mM |
| HEPES KOH (pH=8) | 55 | mM |
| Magnesium acetate | 14 | mM |
| Potassium acetate | 50 | mM |
| Ammonium acetate | 155 | mM |
| Polyethylene glycol 6000 (PEG) | 3% | (w/v) |
| 3-Phosphoglyceric acid (3-PGA) | 40 | mM |
| Amino acids - mixture I | 2.5 | mM |
| Amino acids - mixture II | 2.5 | mM |
| ATP | 1.2 | mM |
| GTP | 1 | mM |
| UTP | 0.8 | mM |
| IPTG | 1 | mM |
| S30-T7 lysate | 34% | (v/v) |
| DNA | 10 | $\mu g\ ml^{-1}$ |

**Supplementary Table 3**. Synthetic cells' inner solution original composition – before adaptations for production and correct folding of Gaussia luciferase.

| **Reagent** | **final concentration** | |
|---|---|---|
| Sucrose | 200 | mM |
| HEPES KOH (pH=8) | 55 | mM |
| Magnesium acetate | 14 | mM |
| Potassium acetate | 50 | mM |
| Ammonium acetate | 155 | mM |
| Polyethylene glycol 6000 (PEG) | 3% | (w/v) |
| 3-Phosphoglyceric acid (3-PGA) | 40 | mM |
| Amino acids - mixture I | 2.5 | mM |
| Amino acids - mixture II | 2.5 | mM |
| ATP | 1.2 | mM |
| GTP | 1 | mM |
| UTP | 0.8 | mM |

| | | |
|---|---|---|
| IPTG | 1 | mM |
| S30-T7 lysate (**without DTT and β-mercaptoethanol**) | **57%** | **(v/v)** |
| DNA | 10 | µg $ml^{-1}$ |
| **Disulfide bond isomerase C (DsbC)** | **75** | **µg $ml^{-1}$** |
| **Oxidized glutathione (GSSG)** | **4** | **mM** |
| **Reduced glutathione (GSH)** | **1** | **mM** |

**Supplementary Table 4.** Synthetic cells' inner solution composition– after adaptations for production and correct folding of *Gaussia* luciferase. Modifications are highlighted in bold.

| **Reagent** | **final concentration** | |
|---|---|---|
| glucose | 200 | mM |
| HEPES KOH (pH=8) | 55 | mM |
| Magnesium acetate | 14 | mM |
| Potassium acetate | 50 | mM |
| Ammonium acetate | 155 | mM |
| Polyethylene glycol 6000 (PEG) | 3% | (w/v) |
| 3-Phosphoglyceric acid (3-PGA) | 40 | mM |
| Amino acids - mixture I | 2.5 | mM |
| Amino acids - mixture II | 2.5 | mM |
| ATP | 1.2 | mM |
| GTP | 1 | mM |
| UTP | 0.8 | mM |
| IPTG | 1 | mM |
| ultra-pure water | 25% | (v/v) |

**Supplementary Table 5.** Synthetic cells' outer solution composition.

### Supplementary Figures

**Supplementary Figure 1.** Size distribution analysis of 100-nm POPC liposomes using dynamic light scattering (PDI = polydispersity index).

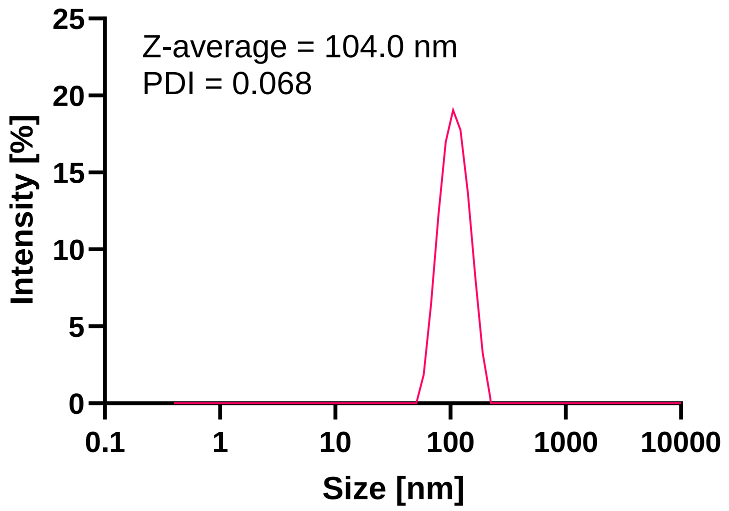

**Supplementary Figure 2.** The effect of cholesterol addition to the liposomal membrane on blue light absorbance in POPC liposomes. The absorbance of 100 nm liposomes with 40 mol% cholesterol or 0 mol% cholesterol at 480 nm was measured. Data is represented as the mean ± standard deviation (n=3 independent samples). Nested two-tailed t-test adjusted P value; *P=0.048.

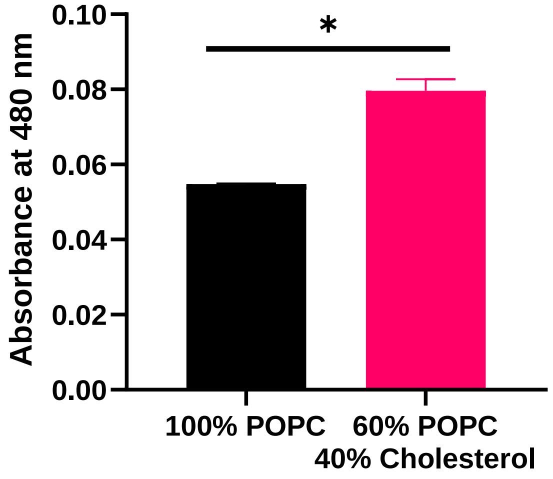

**Supplementary Figure 3**. CryoSEM images distinguishing between synthetic cells and residue oil droplets from the preparation method. Images (a) and (c) were captured with SE2 and InLens imaging detectors. Images (b) and (d) are corresponding images captured with the energy selective backscattered (ESB) detector, used for elemental analysis contrast between organic and aqueous phases. Dark domains in the ESB images represent the organic oil phase and light domains represent the aqueous phase. White arrows indicate synthetic cells and white arrowheads indicate oil droplets.

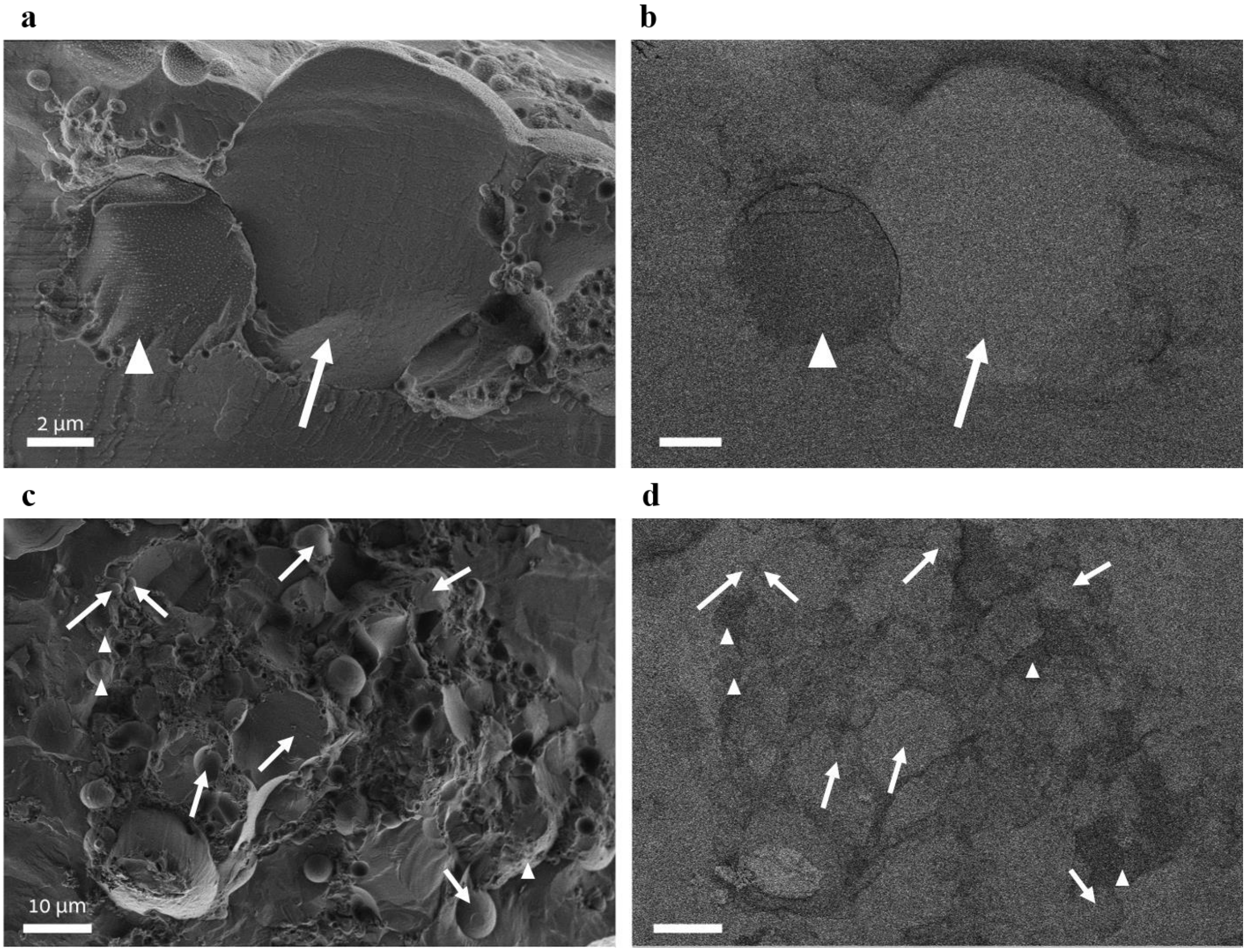

**Supplementary Figure 4.** Absorbance spectrum of liposome formulations of 100 nm liposomes with different lipid membrane compositions.

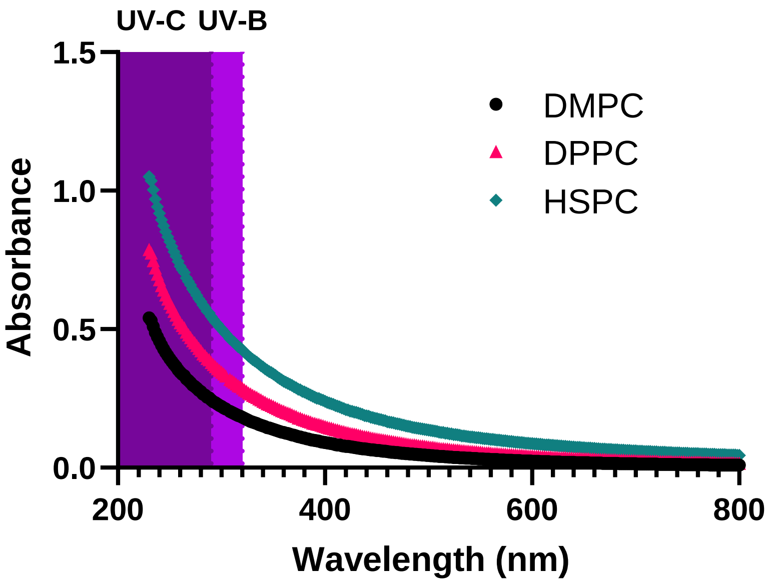

**Supplementary Figure 5.** The effect of different concentrations of oxidized and reduced glutathione on *Gaussia* luciferase production in cell free reactions. Data is represented as the mean ± standard deviation (n=2 independent samples).

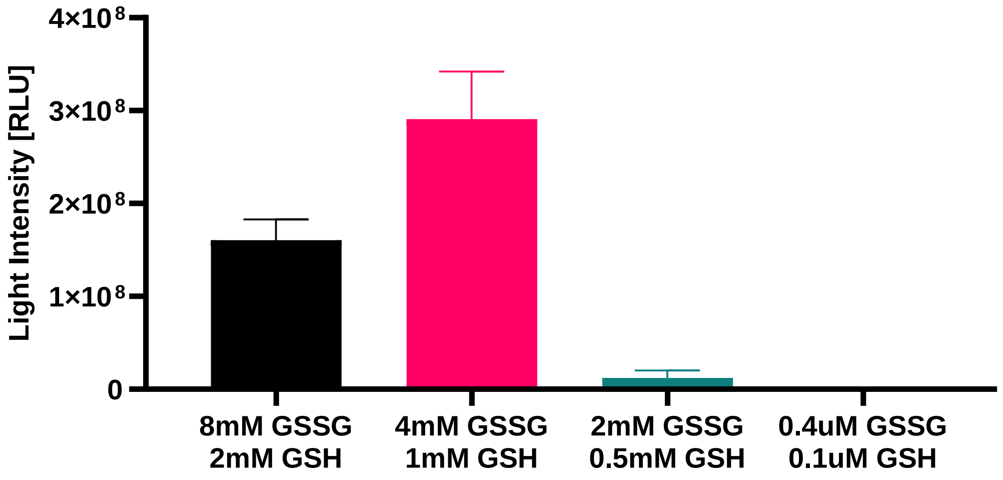

**Supplementary Figure 6.** SDS-Page gel with coomassie blue staining of the purified fraction of disulfide bond isomerase C (DsbC).

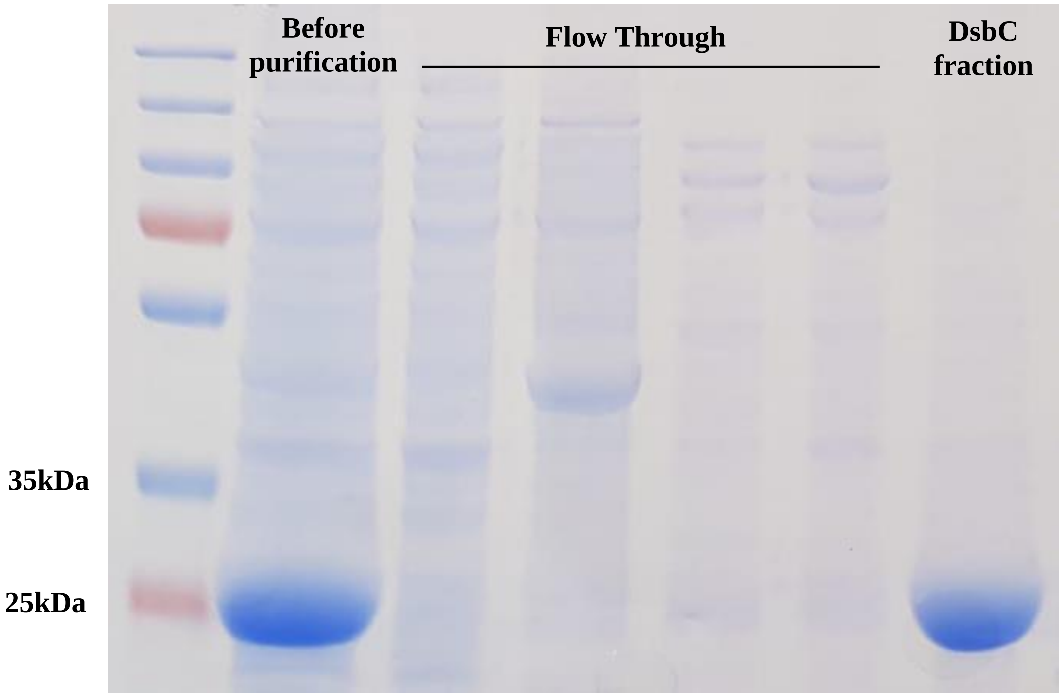

**Supplementary Figure 7.** Light emission from *Gaussia* luciferase expressing synthetic cells with variable concentrations of coelenterazine ranging from 100 μm to 10 nM. Synthetic cells were diluted 400-fold prior to the measurement. Data is represented as the mean ± standard deviation (n=3 independent samples).

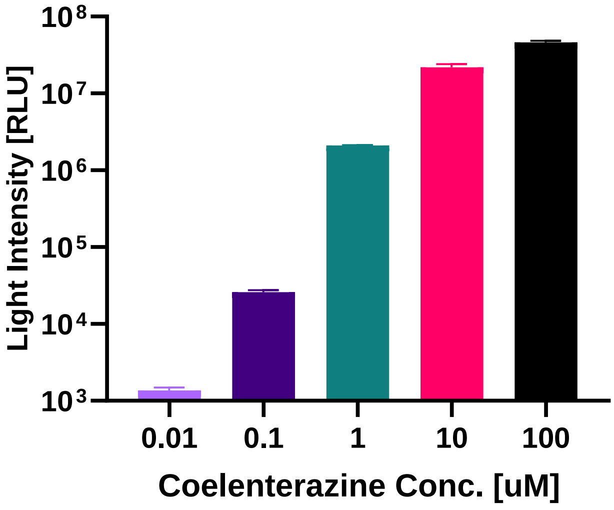

**Supplementary Figure 8.** Induction of photoconidiation in *Trichoderma atroviride* colonies after exposure to blue light (1-minute exposure to 15 mW $cm^{-2}$)

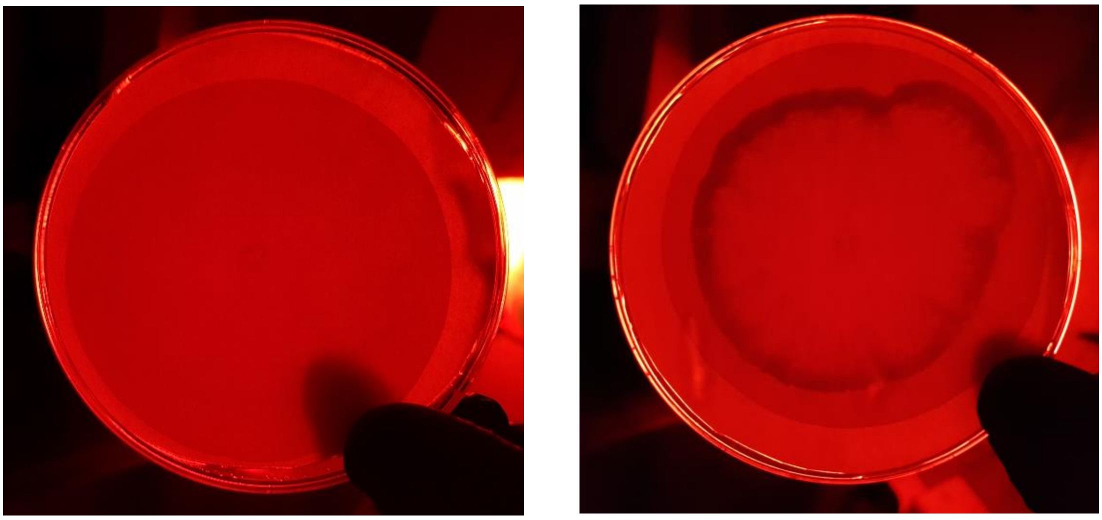

**Incubated in the dark** **Exposed to blue light**

**Supplementary Figure 9.** SDS-Page gel with coomassie blue staining of the purified *Gaussia* luciferase (Gluc) – EL222.

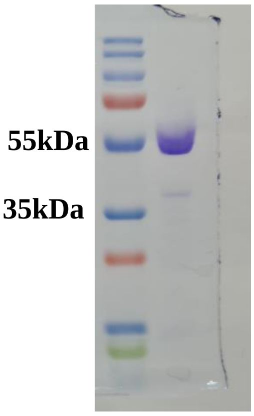

**Supplementary Figure 10.** The effect of coelenterazine addition in different concentrations on the yield of sfGFP in cell free protein synthesis reaction without the *Gaussia* luciferase - EL222 fusion protein. Data is represented as the mean ± standard deviation (n=2 independent samples).

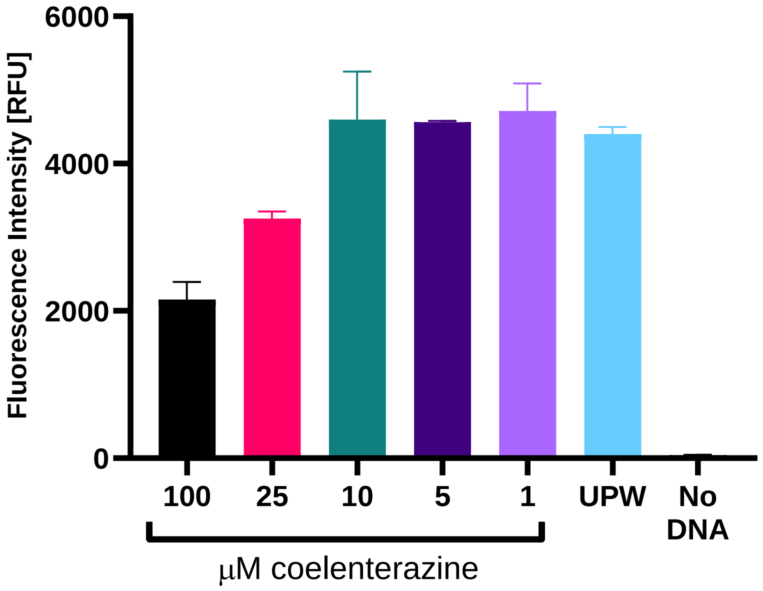

**Supplementary Figure 11.** SDS-Page gel with coomassie blue staining of the his-MBP-mRFP-sspB-Nano after TEV restriction, before and after purification of mRFP-sspB-Nano.

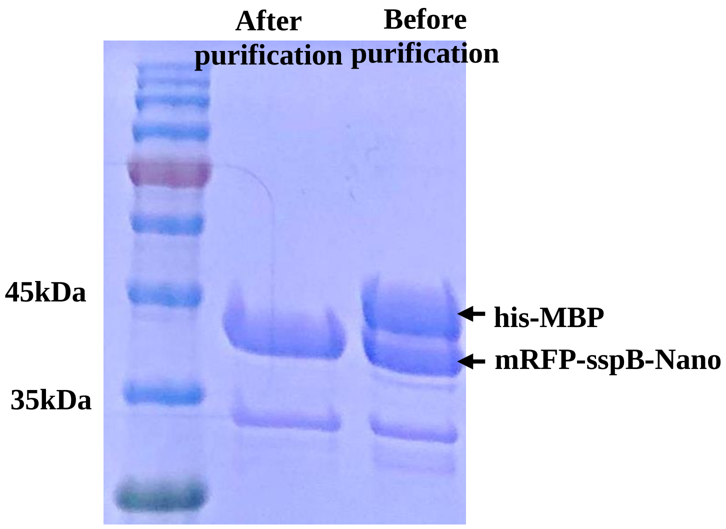

**Supplementary Figure 12.** Recruitment of RFP-sspB-Nano to synthetic cells with iLID functionalized membrane using cell free protein synthesis reactions expressing Gaussia luciferase supplemented with 100 µM coelenterazine. RFP intensity is normalized to the average intensity measured in the dark conditions. SDS-Page gel with coomassie blue staining of the purified Gaussia luciferase (Gluc) – iLID. Data is represented as the mean ± s.e.m. (n=4 independent samples).

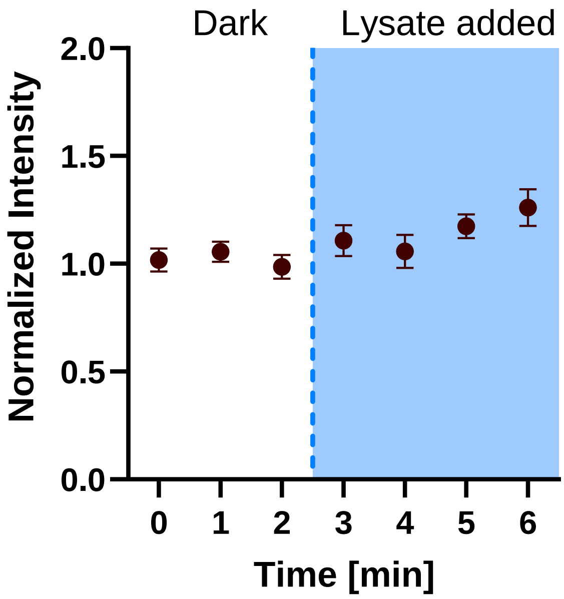

**Supplementary Figure 13.** SDS-Page gel with coomassie blue staining of the purified Gaussia luciferase (Gluc) – iLID.

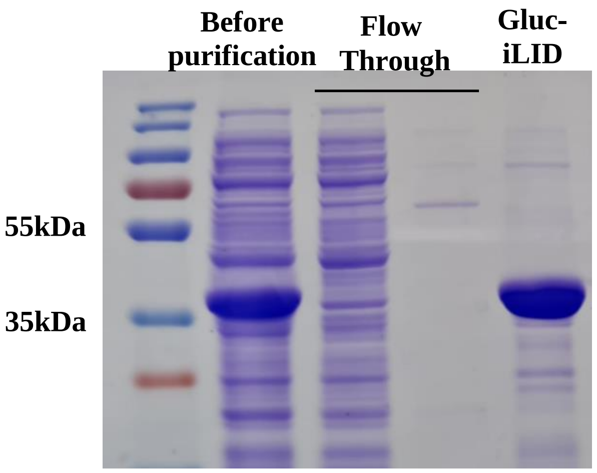

**Supplementary Figure 14.** Recruitment of RFP-sspB-Nano to synthetic cells functionalized with iLID or Gluc-iLID after 4 minutes of exposure to external 488 nm laser light or after four additions of 0.2 nmol of coelenterazine. Intensity is normalized to average RFP intensity in the dark state before exposure to light or substrate. Data is represented as the mean ± standard deviation (n=3 for iLID + coelenterazine, n=10 for iLID + laser and gluc-iLID + laser, n=14 for Gluc-iLID + coelenterazine). Student's two-tailed t-test P values; *P = 0.0135; **P = 0.0092; ****P < 0.0001.

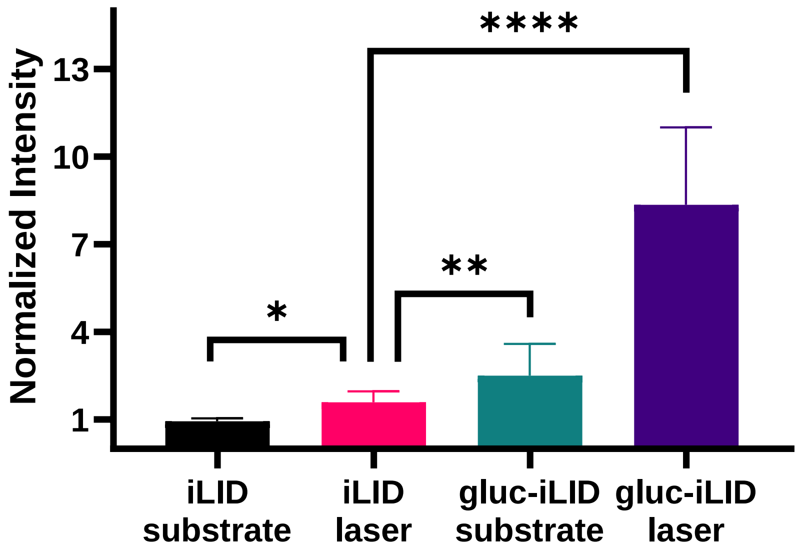
